# Comprehensive Genomic and Evolutionary Analysis of Biofilm Matrix Clusters and Proteins in the *Vibrio* Genus

**DOI:** 10.1101/2024.08.19.608685

**Authors:** Yiyan Yang, Jing Yan, Rich Olson, Xiaofang Jiang

## Abstract

*Vibrio cholerae* pathogens cause cholera, an acute diarrheal disease resulting in significant morbidity and mortality worldwide. Biofilms in vibrios enhance their survival in natural ecosystems and facilitate transmission during cholera outbreaks. Critical components of the biofilm matrix include the *Vibrio* polysaccharides produced by the *vps*-1 and *vps*-2 gene clusters and the biofilm matrix proteins encoded in the *rbm* gene cluster, together comprising the biofilm matrix cluster. However, the biofilm matrix clusters and their evolutionary patterns in other *Vibrio* species remain underexplored. In this study, we systematically investigated the distribution, diversity, and evolution of biofilm matrix clusters and proteins across the *Vibrio* genus. Our findings reveal that these gene clusters are sporadically distributed throughout the genus, even appearing in species phylogenetically distant from *V. cholerae*. Evolutionary analysis of the major biofilm matrix proteins RbmC and Bap1 shows that they are structurally and sequentially related, having undergone structural domain and modular alterations. Additionally, a novel loop-less Bap1 variant was identified, predominantly represented in two phylogenetically distant *Vibrio cholerae* subspecies clades that share specific gene groups associated with the presence or absence of the protein. Furthermore, our analysis revealed that *rbmB*, a gene involved in biofilm dispersal, shares a recent common ancestor with Vibriophage tail proteins, suggesting that phages may mimic host functions to evade biofilm-associated defenses. Our study offers a foundational understanding of the diversity and evolution of biofilm matrix clusters in vibrios, laying the groundwork for future biofilm engineering through genetic modification.

## Introduction

*Vibrio cholerae*, the pathogen responsible for cholera, causes an acute diarrheal disease that can lead to hypotonic shock and death. Annually, it infects 3-5 million people, resulting in 100,000– 120,000 deaths (1). *V. cholerae* forms biofilms—surface-associated communities encased in a matrix—which enhance survival in ecosystems, and transmission during outbreaks (2, 3), while providing protection from environmental stresses like nutrient scarcity, antimicrobial agents, predation by unicellular eukaryotes, and attack by phages (4–6).

The biofilm matrix is primarily comprised of <u>V</u>ibrio <u>p</u>oly<u>s</u>accharide (VPS), making up approximately half of its mass and essential for biofilm 3D structural development (7–9). Genes involved in VPS production are organized into two *vps* gene clusters, *vps*-1 and *vps*-2. A gene cluster in this study is defined as a group of closely located genes on a chromosome that are often functionally related and may include multiple operons. The *vps*-1 gene cluster contains 12 genes (*vpsU*, VC0916 and *vpsA-K*, VC0917-VC0927) while the *vps*-2 gene cluster is relatively shorter only containing 6 genes (*vpsL-Q*, VC0934-VC0939) (7, 9). Meanwhile, biofilm matrix proteins, such as RbmA, RbmC and Bap1, encoded by *rbmA* (VC0928), *rbmC* (VC0930) and *bap1* (VC1888), respectively, are crucial for preserving the structural integrity of the wild-type biofilm (10, 11), among which RbmA and RbmC are encoded in a *rbm* (rugosity and biofilm structure modulator) gene cluster separating the two *vps* gene clusters. The gene encoding Bap1 is distant from the *rbm* gene cluster, yet it also modulates the development of corrugated colonies and is crucial for biofilm formation (11–13). RbmA, as a biofilm scaffolding protein involved in cell-cell and cell-biofilm adhesion, is required for rugose colony formation and biofilm structure integrity in *V. cholerae* (10, 11, 13–15). The other two major biofilm matrix proteins, RbmC and Bap1, are homologues sharing 47% sequence similarity and containing overlapping domains to facilitate their robust adhesion to diverse surfaces (11, 16). Both proteins have a conserved β-propeller domain with eight blades and at least one β-prism domain. RbmC, however, is characterized by two β-prism domains and additional tandem β/γ crystallin domains, known as M1M2 (16, 17). Most notably, Bap1’s β-prism contains an additional 57-amino acid (aa) sequence which promotes *V. cholerae* biofilm adhesion to lipids and abiotic surfaces while RbmC mainly mediates binding to host surfaces through recognition of N- and O-glycans and mucins (16). Another interesting gene in the *rbm* gene cluster is *rbmB* (VC0929), which encodes a putative polysaccharide lyase that has been proposed to have a role in VPS degradation and cell detachment (11, 18–20). Other genes included in the *rbm* gene cluster are *rbmDEF* (VC0931-VC0933). Together, the *vps*-1, *rbm* and *vps*-2 gene clusters comprise a functional genetic module — the *V. cholerae* biofilm matrix cluster (*V. cholerae* BMC or VcBMC) (18).

The biofilm matrix cluster has primarily been investigated in commonly studied *V. cholerae* strains and a few other *Vibrio* species (21–24). However, it has not yet been systematically studied at the strain level within *V. cholerae* or more extensively across the *Vibrio* genus. Since the biofilm matrix cluster encodes proteins for VPS synthesis and matrix proteins, which are the major components of *Vibrio* biofilms, a systematic genomic analysis of this cluster and the identification of relevant genes across the *Vibrio* genus can provide a prospective and comprehensive view of the genetic basis underlying VPS production and biofilm formation.

In this study, we comprehensively annotated the genes involved in the biofilm matrix cluster to explore their distribution, diversity and gene synteny by conducting large-scale comparative genomics and phylogenetic analyses on 6,121 Vibrio genomes spanning 210 species across the entire *Vibrio* genus as well as within the *V. cholerae* species. We observed not only a prevalent presence of this cluster in *V. cholerae* but also in other distantly related species. Our analysis reveals a distinct evolutionary pattern for the *vps*-1 and *vps*-2 gene clusters: genes in the *vps*-2 gene cluster often co-located with *rbmDEF* genes, while *vps*-1 genes are commonly adjacent to *rbmABC* genes. This suggests a functional relatedness between them and explains why these two *vps* gene clusters are separated by a *rbm* gene cluster in contemporary *V. cholerae* strains. Additionally, we inferred that the *bap1* genes originated as an ancient duplication of *rbmC* in a clade of species closely related to *V. cholerae*, while *rbmC* genes are present in two major clades and may have undergone structural domain alterations throughout their evolutionary history. Furthermore, a novel loop-less Bap1 variant was identified, predominantly found in two phylogenetically distant *Vibrio cholerae* subspecies clades that share gene groups linked to the presence/absence of the protein. Finally, our findings suggest that RbmB, a putative VPS degradation enzyme, are evolutionarily related to Vibriophage pectin lyase-like tail proteins. The systematic and accurate curation of biofilm matrix clusters and their proteins not only enhances our understanding of *Vibrio* biofilm formation from a genomic view but also offers insights for developing strategies to engineer and control biofilms.

## Results

### Biofilm matrix clusters are found in phylogenetically distant *Vibrio* species

Leveraging over 6,000 genomes from Genome Taxonomy Database (GTDB r214) (25) across the *Vibrio* genus, we systematically annotated the proteins within the biofilm matrix clusters and depicted an overview of the cluster’s gene occurrences spanning 209 *Vibrio* species and seven *V. cholerae* subspecies (Fig.1A). We defined a full biofilm matrix cluster if it contains the 12 key *vps* genes (namely *vpsAB*, *vpsDEF*, *vpsIJK*, and *vpsLMNO*) whose deletions have been shown to cause a dramatic reduction in VPS production and biofilm formation (9) and all of the *rbm* genes.

**Figure 1.**
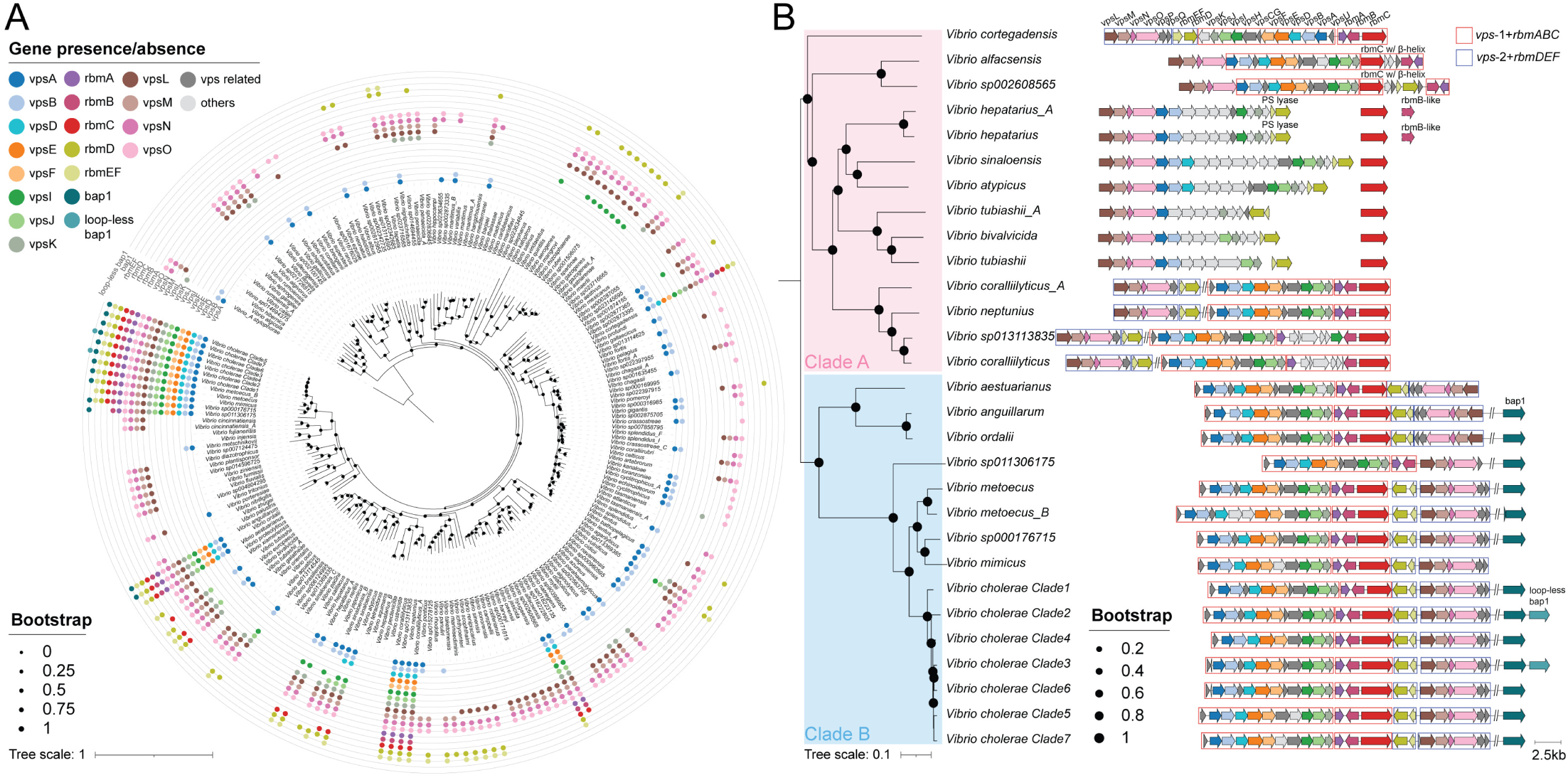
The distribution of biofilm matrix clusters across the *Vibrio* genus. (A) The phylogenomic tree with the presence and absence of important genes in biofilm matrix clusters mapped to tips representing 216 Vibrio (sub)species. The tree was rooted with the representative genome of *Vibrio_A stylophorae* species (NCBI Assembly accession=GCA_921293875.1). (B) Gene syntenies for biofilm matrix clusters in 29 (sub)species that possess biofilm matrix protein encoding genes (*rbmC* and/or *bap1*) are illustrated using the same color palette as in panel A and the phylogenomic tree displayed is a subtree derived from the tree in panel A. The clusters are aligned with each other using the *rbmC* gene as the anchor. Genes that are not concatenated are located on different contigs, whereas genes separated by the “//” symbol are found in the same contig but are hundreds of genes away from each other. The red boxes highlight the proximity of the *vps*-1 and *rbmABC* genes within the genome, while the blue boxes indicate the close genomic location of the *vps*-2 and *rbmDEF* genes. The *rbmE* and *rbmF* genes are combined under the single gene name *rbmEF* due to overlaps in their gene sequences and frequent annotations as a single gene. Similarly, the *vpsC* and *vpsG* genes are merged into one gene name, *vpsCG*, as they both share a highly similar domain. PS: Polysaccharide.

We reconstructed a *Vibrio* species tree, which shares a similar topology to that in a previous study (26), and mapped the presence and absence of the key *vps* genes and *rbm* genes to the tree tips. It is interesting to discover that, using this criterion, the full biofilm matrix clusters not only exist in *V. cholerae* and closely related species (such as *V. metoecus* and *V. mimicus*) but are also sporadically distributed across the *Vibrio* genus in distant species like *V. anguillarum*, *V. ordalii*, *V. aestuarianus*, *V. coralliilyticus*, *V. neptunius* and *V. cortegadensis* (Fig.1A). Among all genes, *vps*-2 genes are the most prevalent genes with *vpsL* existing in 50% of the species, *vpsM* in 41.2%, *vpsN* in 58.3% and *vpsQ* in 64.4% following by *vps*-1 genes *vpsA* (33.3%) and *vpsB* (33.8%). The higher prevalence of *vps*-2 genes is due to the identification of *vps*-2 similar loci in our data, such as the *cps* (capsular polysaccharide) locus in *Vibrio parahaemolyticus*, the *wcr* (capsular and rugose polysaccharide) locus in *Vibrio vulnificus*, and *vps*-2-like loci in *Aliivibrio fischeri*, all of which contain homologs of *vpsLMNO* (Supplementary Figure 1) (27–31). It is important to note that these loci contain genes associated with functions other than VPS production in biofilms, such as capsular polysaccharide synthesis. Therefore, they are less likely to represent true *vps*-2 gene clusters and are instead designated as *vps*-2 similar gene clusters in this study.

We next investigated the gene synteny within the biofilm matrix cluster to gain insights on how the *vps-1*, *vps-2* and *rbm* gene clusters have evolved during the speciation of *Vibrio* species (Figure 1B and Supplementary Figure 2). The Vibrio (sub)species clearly form two major clades, Clades A and B, each of which are featured with different patterns in the biofilm matrix clusters (Fig.1B). The examination of the isolation sources and potential hosts of *Vibrio* species in these clades indicates that Clade A species are primarily isolated from marine water and from healthy or diseased invertebrates such as prawns, corals, and bivalve mollusks like clams and oysters (Supplementary Table 1). In contrast, species in Clade B are mostly found in seawater and brackish waters, inhabiting both invertebrate and vertebrate hosts, including fish (such as *V. aestuarianus*, *V. ordalii*, and *V. anguillarum*) and humans (such as *V. metoecus*, *V. mimicus*, and *V. cholerae*), and often acting as pathogens (Supplementary Table 1).

From Figure 1B, we also observed that *rbmA* genes are absent in seven *Vibrio* species from Clade A (i.e. *V. hepatarius_A*, *V. hepatarius*, *V. sinaloensis*, *V. atypicus*, *V. tubiashii_A*, *V. tubiashii*, and *V. bivalvicida*) despite the presence of *rbmD* and *rbmEF* genes in the same operon and the presence of distant *rbmC* genes. Although these species are phylogenetically distant, we observed conservation in the neighborhoods of their *rbmC* genes. These *rbmC* genes are often immediately adjacent to a gene containing a methyl-accepting chemotaxis domain and are close to an operon encoding a system for the uptake and metabolism of disaccharides, suggesting their potential involvement in sugar binding process (Supplementary Figure 3 and Supplementary Table 2). These species typically possess several, but not all, *vps*-2 similar and *vps*-1 similar genes. For genes not annotated as *vps*-like genes, most of them are glycosyltransferases, acyltransferases and polysaccharide biosynthesis proteins, which might be responsible for the synthesis, modification and export of VPS (Supplementary Figure 2 and Supplementary Table 3).

Additionally, we observed that *vps*-1 gene clusters tend to co-locate with *rbmABC* genes, while *vps*-2 gene clusters consistently pair with *rbmDEF* genes (see red and blue boxes in Figure 1B). This patten is evident in a sub-lineage of Clade A, which includes *V. coralliilyticus, V. coralliilyticus_A*, *V. neptunius*, and *V. sp013113835*. In this sub-lineage, *vps*-2 and *rbmDEF* genes are joined but remain distant from the joined *vps*-1 and *rbmABC* genes (Fig.1B). In contrast, Clade B features an intact biofilm matrix cluster, where the *vps*-2 genes and *rbmDEF* genes are consistently adjacent and linked to the joined *vps*-1 and *rbmABC* genes. We also observed that in *V. aestuarianus*, *V. anguillarum*, and *V. ordalii*, the *vps*-2 gene cluster is in the opposite orientation compared to other species within Clade B. Overall, the consistent co-location of the *vps*-1 genes with *rbmABC* and the *vps*-2 genes with *rbmDEF* in several Clade A species and across the whole Clade B suggests their functional associations. This organization may also help explain the intact biofilm matrix clusters commonly observed in *Vibrio cholerae* strains, where the two *vps* gene clusters, separated by a *rbm* gene cluster, could result from the merging of *rbmABC* and *rbmDEF* genes.

### Biofilm matrix proteins RbmC and Bap1 experienced structural diversification during evolution

RbmC and Bap1, two major biofilm matrix proteins in Vibrio biofilms that share 47% sequence identity, have been shown in previous studies to possess both shared and distinct domains. This suggests that they are functionally and evolutionarily related, with potential domain gain and loss occurring during their evolution (11, 16). Furthermore, we are interested in Bap1-encoded gene due to its higher mutation frequency of 0.0718 compared to all *rbm* genes, implying it may have undergone a stronger positive selection pressure (Supplementary Figure 4). To examine the origin and divergence of these two matrix proteins, we compiled a data set consisting of 2,004 *rbmC* and 2,062 *bap1* genes identified across the *Vibrio* genus.

There are three extra RbmC variants as well as one Bap1 variant (Supplementary Figures 5 and 6). Two of the RbmC variants differ from the standard RbmC protein by having none or only one of the two mucin-binding domains (referred to as M1M2-less and partial M1M2 RbmC, respectively). Most of the M1M2-less RbmC (59%) and partial M1M2 RbmC (85%) proteins were found to have signal peptides, likely still functioning as an intact protein despite losing domains. Five genes from *V. alfacsensis* and *V. sp002608565* genomes represent the third RbmC variant, which show an averaged ∼43% and ∼52% similarity to the standard *rbmC* and *bap1*, respectively. This variant has β-propeller and β-helix domains, with corresponding genes located in positions typically associated with *rbmC* genes in the biofilm matrix clusters. Therefore, they are labeled as “*rbmC* w/ β-helix” (Fig. 1B). On the other hand, *bap1* genes are exclusively found in *V. cholerae* and its closely related species within Clade B. Upon examining the neighboring genes of *bap1*, we identified a duplicate *bap1* gene, that encodes a Bap1 variant, directly adjacent to the standard *bap1* (Fig.1B). It shares all of the domains with standard Bap1 protein but lacks the 57aa loop in the β-prism domain and is therefore referred to as loop-less Bap1 (16). Taken together, we identified a total of six structural groups representing different protein variants: RbmC with β-helix, M1M2-less RbmC, partial M1M2 RbmC, standard RbmC, standard Bap1 and loop-less Bap1 (Fig.2A).

**Figure 2.**
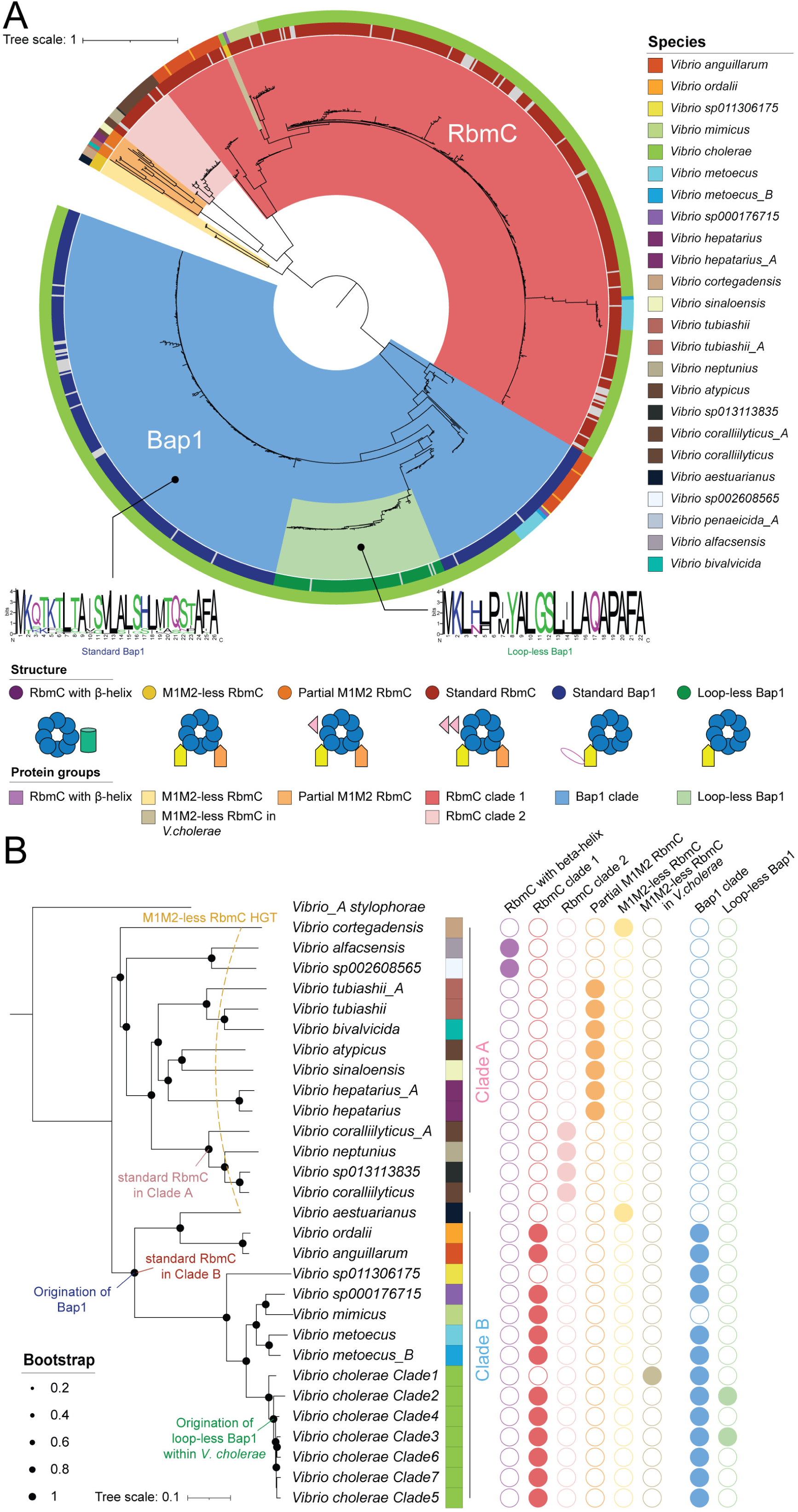
The gene tree and evolutionary analysis for RbmC and Bap1 proteins. (A) The gene tree was built with non-redundant codon sequences of 514 RbmC and 483 Bap1 proteins, which is rooted at the midpoint. The outer circle indicates the species of origin, while the inner circle indicates the protein structural features with grey representing truncated proteins. The cartoons at the bottom demonstrate the domain composition for the corresponding structures. Color ranges indicate different protein groups based on both structural features and phylogenetic relationships, whose legend was put under the corresponding structural features. Note that the RbmC with a β-helix domain was omitted from the gene tree due to it causing a poor multiple sequence alignment. The sequence logos for the signal peptides are shown for the Bap1 clade and loop-less Bap1 clade. (B) The distribution of 9 protein groups along the phylogenetic tree suggests evolutionary events for *rbmC* and *bap1* genes. The tree replicates the one in Fig.1B while retaining the outgroup species. The species and protein group colors are consistent with those in panel A.

Next, after sequence redundancy removal, a codon-based phylogenetic tree was constructed. The phylogeny indicates that the RbmC and Bap1 form two distinct clades, and the long branch connecting them suggests their distant divergence. Protein sequences from the same structural group typically cluster together, although there are exceptions. For instance, a group of genes encoding M1M2-less RbmC is exclusively found in *V. cholerae* and nested within the largest standard RbmC clade, while genes for loop-less Bap1 fall into a subclade within the standard Bap1 clade (Fig.2A). Taking this phylogenetic information into consideration, we have further divided all of the protein sequences into eight protein groups: RbmC with β-helix, M1M2-less RbmC, M1M2-less RbmC in *V. cholerae*, partial M1M2 RbmC, RbmC clade 1, RbmC clade 2, Bap1 clade, and loop-less Bap1 (see gene group cartoon illustrations in Fig.2A).

We mapped these protein groups onto the *Vibrio* species tree tips to infer their evolutionary events. The eight protein groups demonstrated distinct patterns between Clades A and B (Fig.2B). Genes encoding RbmC variants are observed across the species in Clade A, but no Bap1 encoded genes are found. We also observed that RbmC has undergone a series of alterations in the M1M2 domains with a β-helix domain replacing the original M1M2 and β-prisms domains in Clade A. Genes encoding standard RbmC are prevalent in Clade B, in contrast to their restricted presence in a subclade of Clade A (see nodes labeled as “standard RbmC in Clade A” and “standard RbmC in Clade B” in Fig.2B). Genes encoding Bap1 are also found exclusively in Clade B, suggesting that Bap1 genes originated at the ancestral node of this clade (see node labeled as “Origination of Bap1” in Fig.2B). The phylogenetic analysis of the β-propeller domains suggests that Bap1 may have diverged from the ancestor of standard RbmC in both Clade A and Clade B (Supplementary Figure 7). It has been reported that the sequence of Bap1’s β-prism diverges from the β-prisms in RbmC (17), and our analysis further shows that Bap1’s β-prism domains are closer to RbmC’s first β-prism domain (β-prism C1) than to the second (β-prism C2), sharing the most recent common ancestor with RbmC’s first β-prism domains exclusively in Clade A (Supplementary Figure 8). This observation aligns with previous findings (14, 16). In addition, the genes encoding loop-less Bap1 are likely to originate from a *V. cholerae* lineage within Clade B (see node labels as “Origination of loop-less Bap1 within *V. cholerae*” in Fig.2B).

A horizontal gene transfer event (HGT) of genes encoding M1M2-less RbmC was observed from *V. cortegadensis* species in Clade B to *V. aestuarianus* species in Clade A. We inferred this to be a result of horizontal gene transfer (see yellow dashed line in Fig.2B) because the genes encoding M1M2-less RbmC, while phylogenetically closest (Fig.2A), are found in two distantly related species in the *Vibrio* species tree (Fig.2B). Interestingly, the biofilm matrix clusters in the genomes of these two species are also highly similar and only slightly differ in the direction and location of the *rbmABC* genes (Fig.1B). In terms of the absence of M1M2 domains in RbmC proteins within *Vibrio cholerae* Clade 1, it is likely the result of a domain loss event in the standard RbmC proteins. This is supported by them forming into a distinct subclade within the standard RbmC Clade 1 on the gene tree (Fig. 2A).

### Loop-less Bap1 proteins are predominantly found in two distant subspecies clades of *V. cholerae*

The standard Bap1 protein and the loop-less variant are predicted to be highly similar in both structures (TM-score=0.8020) and sequences (identity=78.5%) (Supplementary Figure 5E-F). In addition, a 22aa signal peptide was predicted at the N-terminus of loop-less Bap1, which differs in sequence pattern and peptide length from that of the standard Bap1, whose signal peptide is 26aa (Fig.2A). Therefore, despite the absence of a loop and the likely loss of adhesion ability (16), the presence of a signal peptide in the loop-less Bap1 suggests that the protein is still likely to be expressed and secreted.

We next examined the distribution of loop-less Bap1 in the *V. cholerae* subspecies phylogenetic tree. The phylogeny reveals that *V. cholerae* is divided into seven distinct subspecies clades, with loop-less Bap1-encoding genes predominantly enriched in Clades 2 and 3, and a few scattered in Clade 5 (Fig.3). Clades 2 and 3 are phylogenetically distant (Fig.3A), suggesting that the predominant presence of loop-less Bap1 in these clades may reflect selective pressure acting on the protein. While all strains in Clade 2 are classified as *Vibrio cholerae* by GTDB, 38.75% are identified as *Vibrio paracholerae* according to NCBI taxonomy (noting that GTDB does not recognize *Vibrio paracholerae* as a species), indicating its close relationship to *V. paracholerae*. In Clade 3, no clear taxonomic patterns were observed, though the strain *Vibrio albensis* ATCC 14547 (reclassified as *V. cholerae* species in GTDB) is included. Clade 5 is noteworthy, as it includes strains associated with 7PET (7^th^ pandemic El Tor) or a putative 7PET lineage (Supplementary Figure 9 and Supplementary Table 4).

**Figure 3.**
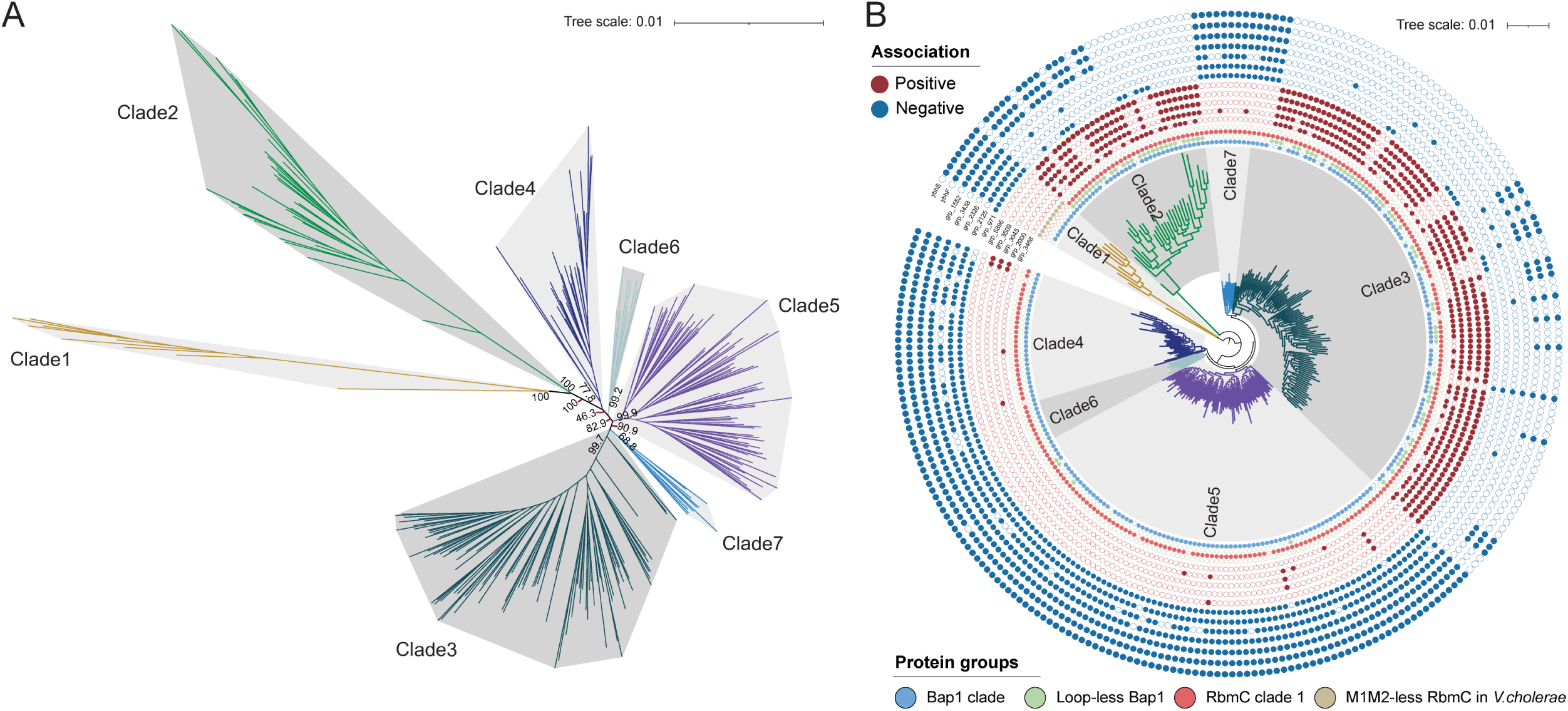
Loop-less Bap1 encoded genes are predominantly found in two distant *V. cholerae* clades, which share specific gene groups associated with the presence/absence of the protein. (A) Unrooted phylogenomic tree of *V. cholerae* species (N=273), with bootstrap values displayed at clade ancestral nodes and nodes representing clade divergence. (B) The phylogenomic tree for *V. cholerae* species was built with protein sequences from the core genes found by Roary (32). The tree was rooted at Clade 1. The presence and absence of RbmC/Bap1 variants (inner circles, using the same palette in Fig.2) and gene groups either positively (dark red) or negatively (dark blue) associated with loop-less Bap1-positive strains (outer circles) are mapped to the tips.

To further investigate the genomic traits and potential roles in metabolic pathways of strains harboring loop-less Bap1, we conducted a comparative genomic analysis using Evolink (33) (see details in Methods), which identified gene groups predominantly present and absent in loop-less Bap1-positive strains, referred to as positively and negatively associated gene groups, respectively. We identified five positively and seven negatively associated gene groups (Fig.3B and Supplementary Table 5). Among the positively associated groups, group_3468 is annotated as putative diguanylate cyclase (DGC) with a GGDEF domain and located close to methyl-accepting chemotaxis-related proteins (Supplementary Figure 10A). Among the negatively associated groups, group_1552 encodes an HlyD family secretion protein and is part of *ybhGFSR* system with other two negatively associated gene groups, *ybhF* and *ybhS* (34). Group_2125 contains a methyl-accepting chemotaxis protein signaling domain and is adjacent to a gene encoding a chitinase (*chiA*) (Supplementary Figure 10B). Despite having opposing associations, group_3045 and group_971, are both predicted to function as histidine kinases involved in signal transduction and positioned near group_525, which is annotated as a c-di-GMP phosphodiesterase (Supplementary Figure 10C). These associations highlight potential regulatory pathways and signaling mechanisms that may influence biofilm formation in loop-less Bap1 positive strains in *V. cholerae*.

### RbmB is evolutionarily related to Vibrio prophage pectin lyase-like tail proteins

We further studied *rbmB* due to its key role in VPS degradation, which regulates biofilm dispersal and cell detachment in *Vibrio cholerae* (11, 19, 20, 35). Its high mutation frequency (0.0604) among all *rbm* genes also suggests a strong positive selection pressure on it (Supplementary Figure 4), highlighting its adaptive significance in promoting biofilm dispersal and its potential as a target for infection control.

By integrating both gene synteny and structural information, we confidently identified 1,760 *rbmB* genes (see details in Methods). We also identified 7,532 genes encoding the pectin lyase-like domain across the Vibrio genus. However, *rbmB* genes account for only 23.4% of these, raising our curiosity about the source and relationships of *rbmB* with other genes. Particularly, given the well-documented role of pectin lyase-like domains in breaking down polysaccharides (36) and their presence in certain phage tail depolymerases that facilitate biofilm degradation (37–39), we explored the possibility that RbmB is evolutionarily related to Vibriophage proteins. To address the abovementioned questions, we constructed a gene tree for Vibrio proteins predicted to have the single-stranded right-handed β-helix (RBH)/pectin lyase-like domains (Fig.4A and Supplementary Figure 11). We observed that over half of the genes (56.1%) are unidentified non-RbmB-encoded genes, and 28.2% are putative pectate lyases. The third largest gene group comprises RbmB-encoded genes (N=319), forming a monophyly in the gene tree (highlighted in red in Fig.4A). The top five species to which these genes belong are *V. cholerae* (N=225), *V. mimicus* (N=20), *V. coralliilyticus* (N=19), *V. metoecus* (N=15) and *V. anguillarum* (N=12) species. Genes in this group have a median length of 408 amino acids and possess signal peptides. This group is closely related to a sister group consisting of 21 non-RbmB-encoded genes (highlighted in yellow in Fig.4A). Together, the two groups are part of a larger clade that includes a large outgroup of 143 non-RbmB-encoded genes (highlighted in blue in Fig.4A). Both groups of 21- and 143-non-RbmB-encoded genes exhibit high structural similarity and moderate sequence similarity to those of the RbmB group, suggesting their close evolutionary relationship (Fig.4B-C).

**Figure 4.**
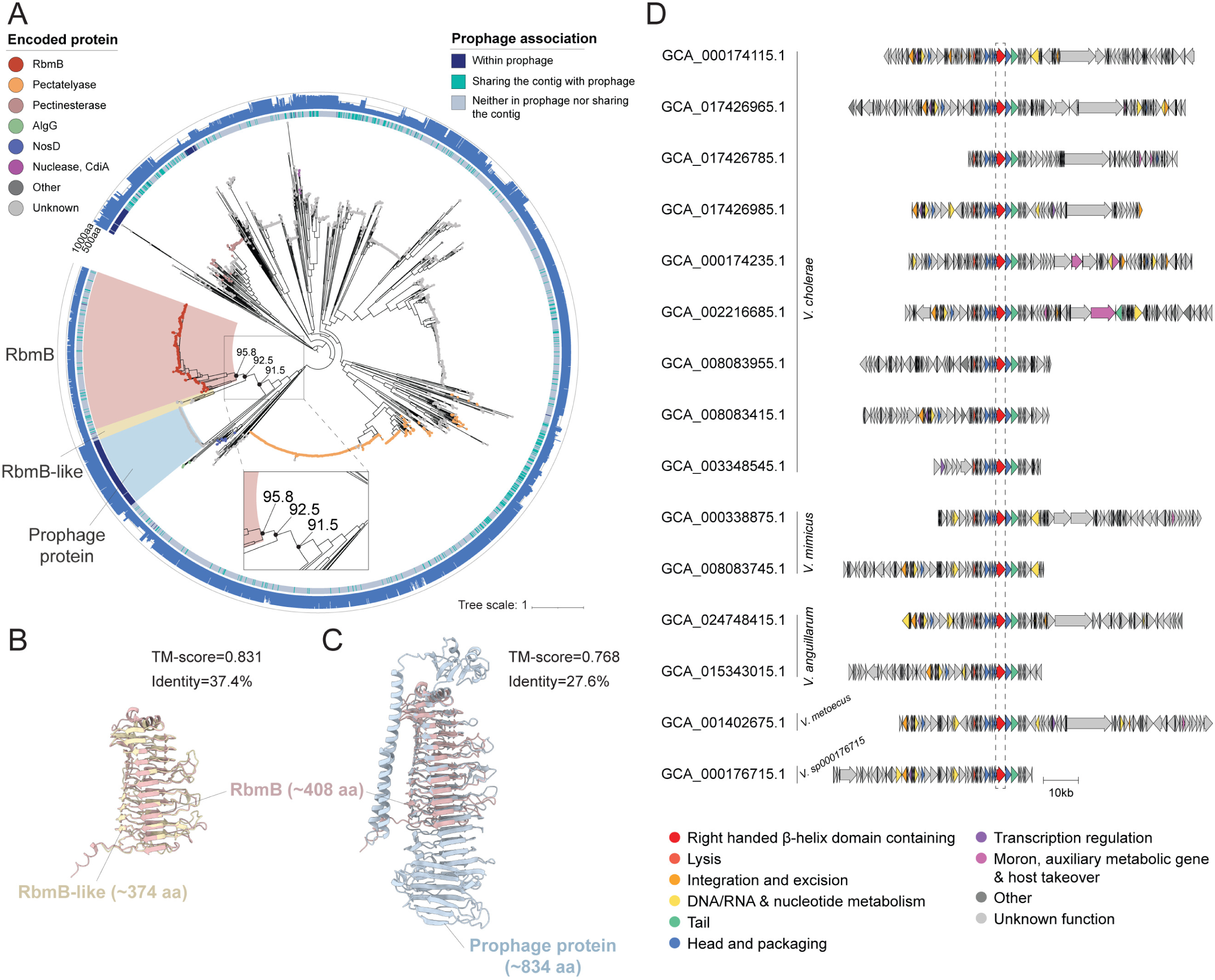
Single-stranded right-handed β-helix (RBH) domain containing gene tree suggests an association between RbmB and prophage proteins. (A) The gene tree was built with non-redundant protein sequences containing single-stranded RBH domains (SUPERFAMILY: SSF51126) and was rooted at the midpoint. Encoded proteins are annotated as colored dots at tips. The inner circle represents the associations of the genes with the prophages found in the same contigs, while the outer circle represents the gene lengths. Bootstrap values are shown at three key internal nodes. The color ranges highlight the clades for RbmB encoded genes (red), RbmB-like encoded genes (yellow) and prophage-related genes (blue). (B-C) Pairwise superimposition of predicted protein structures. The structures displayed are for RbmB (colored red, gene accession: GCA_013111535.1_02619), RbmB-like (colored yellow, gene accession: GCA_002284395.1_03257), and prophage proteins (colored blue, gene accession: GCA_002097735.1_02038). The signal peptides were removed from RbmB and RbmB-like proteins and the structures were predicted by AlphaFold3 (40). (D) Gene syntenies for the 15 representative prophages that possess single-stranded RBH domain containing genes. Each gene synteny is accompanied by the genome accessions from which the prophage fragment was found. Genes encoding the single-stranded RBH domain are colored red, while other genes are colored according to phage functional categories. AlgG: Mannuronan C5-epimerase; NosD: Putative ABC transporter binding protein.

The 21 non-RbmB encoded genes belong to *V. cholerae* (N=9), *V. anguillarum* (N=6), *V. hepatarius* (N=2), *V. hepatarius_A* (N=2) and *V. mimicus* (N=2) species, with a median gene length of 374 amino acids and possessing signal peptides. Thirteen out of the 21 genomes containing these genes also host confidently curated *rbmB* genes, located hundreds of genes away, and all of these genomes additionally contains *rbmC* genes. Taken together, we believe that these genes encode secretory proteins that are functionally different from the real *rbmB* and are named *rbmB*-like genes in this study. The 21 genes likely play distinct roles across different species due to their involvement in varying gene contexts (see *rbmB*-like genes in Fig.1B and Supplementary Table 6). Interestingly, while *V. hepatarius* and *V. hepatarius_A* species lack true *rbmB* genes, they possess putative polysaccharide lyases with β-jelly roll domains located near *vps*-2 similar genes, which might serve as RbmB alternatives for polysaccharide degradation or biofilm dispersal (Supplementary Figure 2 and Supplementary Table 6).

On the other hand, the majority of the 143 non-RbmB encoded genes are from *V. cholerae* (N=124), while the remaining are from *V. mimicus* (N=8), *V. anguillarum* (N=6), *V. metoecus* (N=4) and *V. sp000176715* (N=1) species, with a median gene length of 834 amino acids and lacking signal peptides. One hundred and twenty-six of the 143 genomes containing these genes possess confidently curated *rbmB* genes, which are far from these genes, and all the genomes, except for one, also host *rbmC* or *bap1* genes. Strikingly, we found that 142 of 143 the genes are in the prophage regions. For the only gene not detected in any prophage region in the same contig, it is likely due to the fact that it is the sole gene in the contig that is a relatively short contig that is only 2,667 base pairs long. Gene synteny analysis demonstrated the similarity in the locations of the genes in the 15 representative prophage genomes, where they are situated between two head and packing function-related genes and close to a tail protein (Fig.4D). In addition, BLASTp results showed that all of the 143 genes’ best hits (41) share around 30% identity with the tail fiber protein in Vibrio phage vB_VchM_Kuja (GeneBank accession: MN718199) when queried against the Infrastructure for a PHAge Reference Database (INPHARED, accessed on August 15^th^, 2024) (42), suggesting these genes may also function as part of the phage tail fibers (Supplementary Table 7). Based on the phylogenetic relationships between RbmB, RbmB-like, and prophage pectin lyase-like proteins, we infer that they are derived from a common ancestor, with the prophage proteins diverging before the split of the RbmB and RbmB-like proteins. Overall, our finding marks the first time that RbmB has been demonstrated to be evolutionarily related to Vibriophage pectin lyase-like tail proteins, thus expanding our understanding of their genetic and functional connections.

## Discussion

Bacterial biofilms play a vital role as a lifestyle niche for bacteria in natural environments. They also represent a significant health hazard due to their contribution to persistent infections and the contamination of medical equipment (43–46). Despite their importance in bacterial survival and the challenges they pose in clinical settings, the organization and evolution of the genes encoding the components in biofilm-related clusters have not been extensively studied. A deeper genomic and phylogenetic understanding of these clusters and genes is crucial for the development of innovative genetic engineering strategies that target biofilm-surface interactions and offer alternatives to antibiotic treatments. In this study, using *Vibrio cholerae*—the causative agent of pandemic cholera and a model organism for biofilm studies (18, 47) as well as other related species in the *Vibrio* genus as examples, we propose a framework that integrates comparative genomics, phylogeny, gene synteny analysis and structure prediction to thoroughly characterize biofilm matrix clusters and related proteins, a methodology that can be extended to the study of the biofilm associated clusters and proteins in other bacterial species including important pathogens. This approach has also allowed us to identify domain and modular changes in proteins across their evolutionary timelines, revealing the commonality of domain alterations in *Vibrio* biofilm matrix proteins and their potential implications for biofilm development.

As an alternative to combating antibiotic resistance and biofilm formation in Vibrio pathogens, phage therapies are increasingly attracting attention. Notably, phage host-receptor binding proteins, typically tail fibers or tailspikes, are recognized for their ability to cleave polysaccharides (48–52). Concurrently, *rbmB* genes, encoding RbmB proteins involved in biofilm disassembly, demonstrate significant potential for controlling biofilms and potentially serve as a promising approach to combat Vibrio infections. Interestingly, RbmB proteins and phage tail proteins both feature a common domain—the single-stranded RBH/pectin lyase-like domain—suggesting a potential functional link. However, the evolutionary relationship between these proteins has remained elusive. Here, we reveal that RbmB proteins, along with a group of RbmB-like proteins, share a more recent common ancestor with prophage pectin lyase-like tail proteins than with other pectin lyase-like domain-containing proteins. This suggests that phage tail proteins may be distant homologs of biofilm dispersal proteins, not only highlighting the involvement of phages in biofilm-associated protein evolution but also offering further evidence of phages mimicking host functions to circumvent bacterial defenses. More importantly, the comprehensive annotation of RbmB in *Vibrio* species, combined with insights into Vibrio prophage pectin lyase-like tail proteins, establishes a foundation for a potential biofilm degrader pool. These resources could pave the way for the development of novel protein-based therapies to effectively and precisely target biofilms in emerging Vibrio pathogens.

Our findings clarify many aspects of the *Vibrio* biofilm matrix cluster while also raising new questions. Although we have conducted a comprehensive search for the cluster in the existing genomes across the *Vibrio* genus, it is important to note that this biofilm-associated cluster is VPS-dependent. For *Vibrio* genomes lacking both the *vps*-1 and *vps*-2 genes, it is highly likely that biofilm formation via the clusters curated in this study is not feasible, and these organisms may instead rely on other VPS-independent mechanisms, such as the *syp* loci in *V. parahaemolyticus* and *V. vulnificus* (31). In genomes containing only *vps*-2 similar genes but lacking *vps*-1 genes, it is plausible that the *vps*-2 similar genes are instead integral part of alternative gene operons, such as the *cps* locus in *V. parahaemolyticus* and the *wcr* locus in *V. vulnificus* (31). Therefore, additional VPS-independent biofilm-associated clusters remain to be explored and annotated. For the associated genes identified in loop-less Bap1-positive strains, further experimental validation is required to determine whether they function cooperatively and how they influence bacterial biofilms and behaviors. It is also interesting to explore whether there are polysaccharide lyases or glycosidic hydrolases, aside from RbmB, that could help bacterial cells escape from the biofilm during dispersal. For instance, while RbmB-like proteins are present in *V. hepatarius_A* and *V. hepatarius*, their effectiveness in biofilm disassembly is questionable due to their remote location from other *vps* and *rbm* genes. Instead, polysaccharide lyases containing the β-jelly roll domain may assume this role. It would also be intriguing to uncover how variations in RbmC and Bap1 influence biofilm assembly and to determine the extent to which changes in a single domain or module affect Vibrio phenotypes. We anticipate that these unresolved questions will be addressed through more detailed genomic annotations and experimental studies in the future.

## Methods

### Curation of the biofilm matrix cluster

We downloaded 6,121 genomes classified by GTDB r214 (Genome Taxonomy Database) (25) as Vibrio and Vibrio_A species from NCBI assembly database (53) (accessed on February 18^th^, 2024) (Supplementary Data 1). Genomes were annotated by Prokka v1.14.6 (54) with default parameters. KofamScan (https://github.com/takaram/kofam_scan) (55) and InterProScan v5.63-95.0 (56) (with options “-t p -iprlookup --goterms --pathways” and chunksize of 400) were applied to assign KEGG ortholog and predict domains for the genes with default parameters. These genomes along with their gene protein files (.faa), annotation files (.gff) and kofam annotation files (.kofam.tsv) were used as inputs for ProkFunFind (https://github.com/nlm-irp-jianglab/ProkFunFind) (57) to detect potential biofilm matrix clusters. To prepare the queries for the biofilm matrix protein encoded genes, we have collected a set of KEGG orthologs (i.e. KOfam) covering all *vps* genes as well as the *rbmA* gene from the Kyoto Encyclopedia of Genes and Genomes (KEGG) database (https://www.genome.jp/kegg/) (58). We have also composed a hmm profile for all the *rbm* genes. Any clusters of genes containing more than four of the *vps* or *rbm* genes (with option cluster_min_samples=4) and with a gene neighborhood radius of 18 (with option cluster_eps=18) were assigned a cluster ID as a potential biofilm matrix-associated cluster by ProkFunFind. The 18-gene threshold was determined based on the sum of 12 key *vps* genes and 6 *rbm* genes. The *rbmA*, *rbmB*, *rbmC* and *bap1* as well *vpsE* and *vpsF* genes in an output gene annotation file (.gff) was further recognized and curated in the following section, to generate a refined gene annotation file. The configuration file for ProkFunFind, KOFam list and hmm profile files are provided at https://github.com/nlm-irp-jianglab/ProkFunFind and https://zenodo.org/doi/10.5281/zenodo.11509588. The refined gene annotation output obtained from ProkFunFind is available in Supplementary Data 2.

### Curation and classification of the biofilm matrix proteins RbmC and Bap1

Since Bap1 shares over 40% sequence identity with RbmC, traditional sequence-based computational approaches often perform poorly to distinguish them. Furthermore, these two proteins are usually annotated as hemolysin-like proteins by the NCBI genome annotation pipeline, yet they only share less than 40% identity in the single β-prism domain with hemolysins. Another example lies in the initial scanning of ProkFunFind where both *rbmC* and *bap1* genes have been identified as *rbmC* using hmm profile-based search. Nevertheless, RbmC and Bap1 consist of well-studied domains, which inspires us to leverage structural information to distinguish them. First, 4,066 potential RbmC and Bap1 encoded sequences were obtained by querying WP_000200580.1 (RbmC) and WP_001881639.1 (Bap1) against all protein sequences in *Vibrio* genomes using BLASTp v2.15.0+ (41), with a criteria of > 40% identity, > 250 bit score, and > 200 amino acids in aligned length. Next, to better perform a multiple sequence alignment (MSA), after removing sequence redundancy we excluded the five RbmCs with β-helix encoded genes and only selected high-quality RbmC and Bap1 encoded genes. High-quality genes are those with ≥ 80% identity with a Bap1 query and ranging from 650-700aa in length or with ≥ 80% identity with a RbmC query and ranging from 950-1000aa in length, both with bit scores > 900, while the remaining are classified as low-quality genes. We applied MAFFT v7.475 (59) to align high-quality protein sequences with options “--maxiterate 1000 -- localpair” and aligned low-quality protein sequences by adding them to the previously aligned high-quality genes using MAFFT with option “-add”. The aligned protein sequences were mapped back to the nucleotide sequences to align by codons using PAL2NAL v14 (60). Finally, a codon-based phylogenetic tree was built with the aligned nucleotide sequences using RAxML v8.2.12 (61) by providing a partition file (“-m GTRGAMMA -q dna12_3.partition.txt”), based on which the encoded genes were initially classified as RbmC or Bap1. The detailed structural classification was performed according to the presence and absence of domains in both sequences and structures (Supplementary Data 3-4). The domain boundaries were manually determined by investigating the MSA in Geneious Prime v2023.1.2 (https://www.geneious.com) and double checked with the predicted structures obtained from ESMfold v2.0.0 (62) (Supplementary Data 5). All gene syntenies were annotated using Clinker v0.0.28 (63).

### Curation of RbmB, RbmA, VpsE and VspF proteins

We composed a confident set of *rbmB* genes by first including any genes within a seven-gene distance of either a curated *rbmC* or a putative *rbmA* gene that possess a single-stranded RBH domain (SUPERFAMILY: SSF51126) or are annotated as *rbmB* by a hidden Markov model (HMM) search. The gene distance threshold of seven was determined based on our observation of the maximum number of genes located between *rbmB* and *rbmA* in the current data. Since *rbmA* genes haven’t been thoroughly curated, the neighboring *vps* and *rbm* genes of identified *rbmB* genes adjacent only to a putative *rbmA* gene were manually reviewed to determine if they are real *rbmB* genes. Additionally, ten *rbmB* genes were added to the set because they share over 60% sequence identity and cover more than 90% of the alignment with *rbmB* genes in the confident set. The gene context and the presence of *rbmC* in the same genomes were examined to support the likelihood that these genes are real *rbmB* genes but are not connected to other *rbm* genes due to poor genome assembly and sequencing quality.

Likewise, we curated genes as *rbmA* genes if they are within an eight-gene distance of either a curated *rbmB* or a curated *rbmC* gene, as confirmed in previous sections, that possess two fibronectin type III domains (Gene3D: 2.60.40.3880) or are annotated as *rbmA* by hidden Markov model (HMM) search. The gene distance threshold of eight was determined based on our observation of the maximum number of genes located between *rbmA* and *rbmC* in the current data. For genes located distantly from any *rbmB* or *rbmC* genes but having two fibronectin type III domains, we only included them to the *rbmA* gene set if they, as well as the *rbmB* or *rbmC* genes in the same genomes, are on the edge of contigs, indicating a break in the contig. Regarding genes possessing fewer than two fibronectin type III domains but close to a *rbmB* or *rbmC*, we annotated them as *rbmA* only if they are split into multiple smaller genes or fragmented due to poor genome assembly.

We have cautiously annotated *vpsE* and *vpsF*, as they encode the Wzy-polymerase (VpsE) and Wzx-flippase (VpsF) in the *vps*-1 cluster (64), indicating their important roles in the Wzy/Wzx-dependent VPS synthesis pathway. Any genes within a *vps* gene context that are predicted to be polysaccharide biosynthesis proteins (Pfam: PF13440) and have a polysaccharide biosynthesis C-terminal domain (Pfam: PF14667) or are identified as VpsF family polysaccharide biosynthesis proteins (NCBIfam: NF038256), are regarded as *vpsE* or *vpsF*, respectively. Split and fragmented genes, which only have part or none of the domains, were manually annotated and added if they are close to a well-annotated *vpsF*/*vpsE*.

The gene sequences and typing information in this section are provided as Supplementary Data 6-9.

### Calculating mutation frequency

Given a MSA of nucleotide sequences with N sequences and L positions, the mutations at position *i* are:

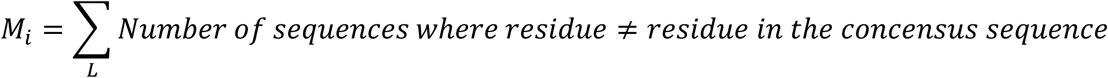

This includes any substitution or gap (“-”) that is not the same as the reference residue.

The mutation frequency is defined as the proportion of mutations relative to all possible positions:

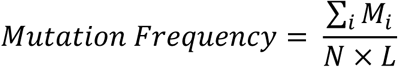

### Selection of *Vibrio* species representative genomes

We didn’t simply use the GTDB representative genomes for the 210 *Vibrio* species in this study. Although the representative genomes generally have high completeness and low contamination, they might have fragmented biofilm matrix clusters and don’t necessarily have the matrix proteins due to genome assembly issues. To take this into consideration, we developed a strategy to pick representative genomes which have maximally reflected the biofilm matrix cluster status at the *Vibrio* species levels. For the 23 species whose genomes possess *rbmC* and/or *bap1* genes, we manually selected the representative genomes that have the most intact biofilm matrix proteins as well as the untruncated RbmC/Bap1 proteins and are representative of the gene synteny of the biofilm matrix cluster in the species. For 73 species in which no biofilm matrix cluster associated proteins was detected, their GTDB representative genomes were used. For the remaining 114 species, 76 of them have multiple genomes. We ranked the genomes in each species higher if they have 1) fewer contigs, implying they have less fragmented contigs, 2) more key *vps*-1 and *vps*-2 genes in the same gene cluster, and 3) more curated *rbm* or *bap1* genes. The genomes meeting these criteria best were selected as the representatives, while the genomes in the 38 single-genome species were picked as species representatives. The final 216 representative genomes for *Vibrio* species and *V. cholerae* subspecies are provided as Supplementary Data 10.

### Pan-genome analysis of *Vibrio cholerae*

A total of 194 core genes were detected and aligned in 1663 *V. cholerae* genomes by pan-genome analysis using the Roary v3.13.0 with options “-i 90 -cd 90 -g 500000 -s -e --mafft” (32). The core gene alignment of a subset of 273 representative genomes with completeness > 90% and contamination < 5% was leveraged to build a phylogenomic tree using FastTree v2.1.11 with default options (65) (Supplementary Data 11). The seven clade representative genomes within V. cholerae species, which have intact biofilm matrix clusters and rbmC/bap1 genes as well as fewer contigs and larger genome lengths, were manually picked for the corresponding clades. The 7PET (7th pandemic El Tor) and putative 7PET lineages were identified by calculating the genomic distance and detecting marker genes with the refence (N16961) using “is-it-7pet” tool (https://github.com/amberjoybarton/is-it-7pet).

### Construction of phylogenomic *Vibrio* species tree

We applied PIRATE v1.0.5 to the 209 *Vibrio* species representative genomes (excluding *V. cholerae*) and seven *V. cholerae* subspecies representative genomes to obtain genus-wise marker genes (with options “-k ‘--diamond’”) (66). PIRATE can rapidly create pangenomes from coding sequences over a wide range of amino acid identity thresholds, thus recognizing the most robust set of core genes. The core gene nucleotide alignment provided by PIRATE was used to build the *Vibrio* species tree using FastTree v2.1.11 with options “-gtr -nt” (Supplementary Data 12). According to GTDB, the *Vibrio_A* genus is more distantly related to the *Vibrio* genus and can serve as a reference group for determining the evolutionary relationships within the *Vibrio* genus. Consequently, the genome of *Vibrio_A stylophorae* was selected as the outgroup to root the tree.

### Identification of loop-less Bap1 positive strains associated gene groups

Given the *V. cholerae* phylogenetic tree, the presence and absence of the gene groups defined by Roary (Supplementary Data 13) and the existence of loop-less Bap1 in genomes, we ran Evolink with default parameters (https://github.com/nlm-irp-jianglab/Evolink) to find five positively and seven negatively associated gene groups related to loop-less Bap1 presence. Evolink is a method for rapid identification of associated genotypes provided a trait of interest and uses phylogenetic approaches to adjust for the population structure in microbial data (33). To confirm that the associated genes identified could be reproduced using alternative methods, we repeated the association analysis with Pyseer using a linear mixed model (67). Pyseer identified 592 significantly associated genes (adjusted p-value < 0.05), 11 of which overlapped with the 12 genes identified by Evolink. The only exception was group_2326, which had an adjusted p-value of 0.108.

### Signal peptide detection

Signal peptides were predicted for RbmC and Bap1-related proteins using SignalP6.0 server (https://services.healthtech.dtu.dk/services/SignalP-6.0/) (68). The signal peptides were aligned with MAFFT v7.475 (59) and visualized as sequence logo using WebLogo server (https://weblogo.berkeley.edu/logo.cgi) (69) (Supplementary Data 14).

### Construction of gene and domain trees

After removing sequence redundancy, single-stranded RBH domain containing protein sequences were aligned using MAFFT-DASH (70) to take structural alignment into consideration. The multiple sequence alignment was next trimmed using TrimAl v1.2rev59 (71) with the -gt 0.2 option to obtain cleaner alignment and used to reconstruct their phylogeny using FastTree v2.1.11 with default options (65).

The β-propeller and β-prism domains sequences were extracted based on domain segmentation of RbmC and Bap1 proteins. The alignment using MAFFT v7.475 (59) were used to build trees using FastTree v2.1.11 with default options (65). All trees were visualized and annotated with iTOL v6 server (https://itol.embl.de/) (72).

The tree files were provided as Supplementary Data 15-17.

### Prophage regions identification

Prophage regions in genomes were detected using VirSorter v2.2.4 (73) with options “--min-length 1000” (Supplementary Data 18). Phage genes within the determined prophage regions were annotated and categorized using Pharokka v1.3.2 (74).

## Data and code availability

The data underlying this article can be accessed through Zenodo (https://zenodo.org/doi/10.5281/zenodo.11509588). All scripts utilized throughout the publication can be accessed through the main branch on the GitHub repository (https://github.com/YiyanYang0728/Vibrio_biofilm_matrix_cluster).

## Acknowledgments

YY and X.J. are supported by the Intramural Research Program of the National Library of Medicine (NLM), National Institutes of Health. This work utilized the computational resources of the NIH HPC Biowulf cluster. (http://hpc.nih.gov).

## Conflicts of interest

The authors declare that there are no conflicts of interest.

## Notes

### Competing Interest Statement

The authors have declared no competing interest.

### Summary of Updates

Section on loop-less Bap1 positive strains associated genes updated to have been deleted or carefully revised and relocated to Discussion section to ensure a clear separation between observed data and speculation, preventing any overstatements; Figure 3 revised; Supplemental figures updated.

## Reference

1. Charles RC, Ryan ET. 2011. Cholera in the 21st century: Current Opinion in Infectious Diseases 24:472–477.

2. Donlan RM, Costerton JW. 2002. Biofilms: Survival Mechanisms of Clinically Relevant Microorganisms. Clin Microbiol Rev 15:167–193.

3. Colwell RR, Huq A, Islam MS, Aziz KMA, Yunus M, Khan NH, Mahmud A, Sack RB, Nair GB, Chakraborty J, Sack DA, Russek-Cohen E. 2003. Reduction of cholera in Bangladeshi villages by simple filtration. Proc Natl Acad Sci USA 100:1051–1055.

4. Gupta P, Mankere B, Chekkoora Keloth S, Tuteja U, Pandey P, Chelvam KT. 2018. Increased antibiotic resistance exhibited by the biofilm of Vibrio cholerae O139. Journal of Antimicrobial Chemotherapy 73:1841–1847.

5. Matz C, McDougald D, Moreno AM, Yung PY, Yildiz FH, Kjelleberg S. 2005. Biofilm formation and phenotypic variation enhance predation-driven persistence of *Vibrio cholerae*. Proc Natl Acad Sci USA 102:16819–16824.

6. Beyhan S, Yildiz FH. 2007. Smooth to rugose phase variation in *Vibrio cholerae* can be mediated by a single nucleotide change that targets c-di-GMP signalling pathway. Molecular Microbiology 63:995–1007.

7. Yildiz FH, Schoolnik GK. 1999. *Vibrio cholerae* O1 El Tor: Identification of a gene cluster required for the rugose colony type, exopolysaccharide production, chlorine resistance, and biofilm formation. Proc Natl Acad Sci USA 96:4028–4033.

8. Yildiz F, Fong J, Sadovskaya I, Grard T, Vinogradov E. 2014. Structural Characterization of the Extracellular Polysaccharide from Vibrio cholerae O1 El-Tor. PLoS ONE 9:e86751.

9. Fong JCN, Syed KA, Klose KE, Yildiz FH. 2010. Role of Vibrio polysaccharide (vps) genes in VPS production, biofilm formation and Vibrio cholerae pathogenesis. Microbiology 156:2757–2769.

10. Fong JCN, Karplus K, Schoolnik GK, Yildiz FH. 2006. Identification and Characterization of RbmA, a Novel Protein Required for the Development of Rugose Colony Morphology and Biofilm Structure in *Vibrio cholerae*. J Bacteriol 188:1049–1059.

11. Fong JCN, Yildiz FH. 2007. The *rbmBCDEF* Gene Cluster Modulates Development of Rugose Colony Morphology and Biofilm Formation in *Vibrio cholerae*. J Bacteriol 189:2319–2330.

12. Moorthy S, Watnick PI. 2005. Identification of novel stage-specific genetic requirements through whole genome transcription profiling of *Vibrio cholerae* biofilm development. Molecular Microbiology 57:1623–1635.

13. Berk V, Fong JCN, Dempsey GT, Develioglu ON, Zhuang X, Liphardt J, Yildiz FH, Chu S. 2012. Molecular Architecture and Assembly Principles of *Vibrio cholerae* Biofilms. Science 337:236–239.

14. Absalon C, Van Dellen K, Watnick PI. 2011. A Communal Bacterial Adhesin Anchors Biofilm and Bystander Cells to Surfaces. PLoS Pathog 7:e1002210.

15. Maestre-Reyna M, Wu W-J, Wang AH-J. 2013. Structural Insights into RbmA, a Biofilm Scaffolding Protein of V. Cholerae. PLoS ONE 8:e82458.

16. Huang X, Nero T, Weerasekera R, Matej KH, Hinbest A, Jiang Z, Lee RF, Wu L, Chak C, Nijjer J, Gibaldi I, Yang H, Gamble N, Ng W-L, Malaker SA, Sumigray K, Olson R, Yan J. 2023. Vibrio cholerae biofilms use modular adhesins with glycan-targeting and nonspecific surface binding domains for colonization. Nat Commun 14:2104.

17. De S, Kaus K, Sinclair S, Case BC, Olson R. 2018. Structural basis of mammalian glycan targeting by Vibrio cholerae cytolysin and biofilm proteins. PLoS Pathog 14:e1006841.

18. Teschler JK, Zamorano-Sánchez D, Utada AS, Warner CJA, Wong GCL, Linington RG, Yildiz FH. 2015. Living in the matrix: assembly and control of Vibrio cholerae biofilms. Nat Rev Microbiol 13:255–268.

19. Bridges AA, Fei C, Bassler BL. 2020. Identification of signaling pathways, matrix-digestion enzymes, and motility components controlling *Vibrio cholerae* biofilm dispersal. Proc Natl Acad Sci USA 117:32639–32647.

20. Díaz-Pascual F, Hartmann R, Lempp M, Vidakovic L, Song B, Jeckel H, Thormann KM, Yildiz FH, Dunkel J, Link H, Nadell CD, Drescher K. 2019. Breakdown of Vibrio cholerae biofilm architecture induced by antibiotics disrupts community barrier function. Nat Microbiol 4:2136–2145.

21. Lilburn TG, Gu J, Cai H, Wang Y. 2010. Comparative genomics of the family Vibrionaceae reveals the wide distribution of genes encoding virulence-associated proteins. BMC Genomics 11:369.

22. Guo Y, Rowe-Magnus DA. 2011. Overlapping and unique contributions of two conserved polysaccharide loci in governing distinct survival phenotypes in *Vibrio vulnificus*. Environmental Microbiology 13:2888–2990.

23. Chodur DM, Rowe-Magnus DA. 2018. Complex Control of a Genomic Island Governing Biofilm and Rugose Colony Development in Vibrio vulnificus. J Bacteriol 200.

24. Gao C, Garren M, Penn K, Fernandez VI, Seymour JR, Thompson JR, Raina J-B, Stocker R. 2021. Coral mucus rapidly induces chemokinesis and genome-wide transcriptional shifts toward early pathogenesis in a bacterial coral pathogen. The ISME Journal 15:3668–3682.

25. Parks DH, Chuvochina M, Rinke C, Mussig AJ, Chaumeil P-A, Hugenholtz P. 2022. GTDB: an ongoing census of bacterial and archaeal diversity through a phylogenetically consistent, rank normalized and complete genome-based taxonomy. Nucleic Acids Research 50:D785–D794.

26. Lin H, Yu M, Wang X, Zhang X-H. 2018. Comparative genomic analysis reveals the evolution and environmental adaptation strategies of vibrios. BMC Genomics 19:135.

27. Smith AB, Siebeling RJ. 2003. Identification of Genetic Loci Required for Capsular Expression in *Vibrio vulnificus*. Infect Immun 71:1091–1097.

28. Güvener ZT, McCarter LL. 2003. Multiple Regulators Control Capsular Polysaccharide Production in *Vibrio parahaemolyticus*. J Bacteriol 185:5431–5441.

29. Grau BL, Henk MC, Garrison KL, Olivier BJ, Schulz RM, O’Reilly KL, Pettis GS. 2008. Further Characterization of *Vibrio vulnificus* Rugose Variants and Identification of a Capsular and Rugose Exopolysaccharide Gene Cluster. Infect Immun 76:1485–1497.

30. Darnell CL, Hussa EA, Visick KL. 2008. The Putative Hybrid Sensor Kinase SypF Coordinates Biofilm Formation in *Vibrio fischeri* by Acting Upstream of Two Response Regulators, SypG and VpsR. J Bacteriol 190:4941–4950.

31. Yildiz FH, Visick KL. 2009. Vibrio biofilms: so much the same yet so different. Trends in Microbiology 17:109–118.

32. Page AJ, Cummins CA, Hunt M, Wong VK, Reuter S, Holden MTG, Fookes M, Falush D, Keane JA, Parkhill J. 2015. Roary: rapid large-scale prokaryote pan genome analysis. Bioinformatics 31:3691–3693.

33. Yang Y, Jiang X. 2023. Evolink: a phylogenetic approach for rapid identification of genotype– phenotype associations in large-scale microbial multispecies data. Bioinformatics 39:btad215.

34. Feng Z, Liu D, Wang L, Wang Y, Zang Z, Liu Z, Song B, Gu L, Fan Z, Yang S, Chen J, Cui Y. 2020. A Putative Efflux Transporter of the ABC Family, YbhFSR, in Escherichia coli Functions in Tetracycline Efflux and Na+(Li+)/H+ Transport. Front Microbiol 11:556.

35. Weerasekera R, Moreau A, Huang X, Nam K-M, Hinbest AJ, Huynh Y, Liu X, Ashwood C, Pepi LE, Paulson E, Cegelski L, Yan J, Olson R. 2024. Vibrio cholerae RbmB is an α-1,4-polysaccharide lyase with biofilm-disrupting activity against Vibrio polysaccharide (VPS). PLoS Pathog 20:e1012750.

36. Burnim AA, Dufault-Thompson K, Jiang X. 2024. The three-sided right-handed β-helix is a versatile fold for glycan interactions. Glycobiology 34:cwae037.

37. Latka A, Drulis-Kawa Z. 2020. Advantages and limitations of microtiter biofilm assays in the model of antibiofilm activity of Klebsiella phage KP34 and its depolymerase. Sci Rep 10:20338.

38. Hughes KA, Sutherland IW, Clark J, Jones MV. 1998. Bacteriophage and associated polysaccharide depolymerases – novel tools for study of bacterial biofilms. Journal of Applied Microbiology 85:583–590.

39. Verma V, Harjai K, Chhibber S. 2010. Structural changes induced by a lytic bacteriophage make ciprofloxacin effective against older biofilm of *Klebsiella pneumoniae*. Biofouling 26:729–737.

40. Abramson J, Adler J, Dunger J, Evans R, Green T, Pritzel A, Ronneberger O, Willmore L, Ballard AJ, Bambrick J, Bodenstein SW, Evans DA, Hung C-C, O’Neill M, Reiman D, Tunyasuvunakool K, Wu Z, Žemgulytė A, Arvaniti E, Beattie C, Bertolli O, Bridgland A, Cherepanov A, Congreve M, Cowen-Rivers AI, Cowie A, Figurnov M, Fuchs FB, Gladman H, Jain R, Khan YA, Low CMR, Perlin K, Potapenko A, Savy P, Singh S, Stecula A, Thillaisundaram A, Tong C, Yakneen S, Zhong ED, Zielinski M, Žídek A, Bapst V, Kohli P, Jaderberg M, Hassabis D, Jumper JM. 2024. Accurate structure prediction of biomolecular interactions with AlphaFold 3. Nature 10.1038/s41586-024-07487-w.

41. Camacho C, Coulouris G, Avagyan V, Ma N, Papadopoulos J, Bealer K, Madden TL. 2009. BLAST+: architecture and applications. BMC Bioinformatics 10:421.

42. Cook R, Brown N, Redgwell T, Rihtman B, Barnes M, Clokie M, Stekel DJ, Hobman J, Jones MA, Millard A. 2021. INfrastructure for a PHAge REference Database: Identification of Large-Scale Biases in the Current Collection of Cultured Phage Genomes. PHAGE 2:214–223.

43. Donlan RM. 2016. Microbial Biofilms, Second Edition. Emerg Infect Dis 22:1142–1142.

44. Hall-Stoodley L, Costerton JW, Stoodley P. 2004. Bacterial biofilms: from the Natural environment to infectious diseases. Nat Rev Microbiol 2:95–108.

45. Costerton JW, Stewart PS, Greenberg EP. 1999. Bacterial Biofilms: A Common Cause of Persistent Infections. Science 284:1318–1322.

46. Flemming H-C, Wingender J, Szewzyk U, Steinberg P, Rice SA, Kjelleberg S. 2016. Biofilms: an emergent form of bacterial life. Nat Rev Microbiol 14:563–575.

47. Nelson EJ, Harris JB, Glenn Morris J, Calderwood SB, Camilli A. 2009. Cholera transmission: the host, pathogen and bacteriophage dynamic. Nat Rev Microbiol 7:693–702.

48. Yen M, Cairns LS, Camilli A. 2017. A cocktail of three virulent bacteriophages prevents Vibrio cholerae infection in animal models. Nat Commun 8:14187.

49. Jensen MA, Faruque SM, Mekalanos JJ, Levin BR. 2006. Modeling the role of bacteriophage in the control of cholera outbreaks. Proc Natl Acad Sci USA 103:4652–4657.

50. Bhandare S, Colom J, Baig A, Ritchie JM, Bukhari H, Shah MA, Sarkar BL, Su J, Wren B, Barrow P, Atterbury RJ. 2019. Reviving Phage Therapy for the Treatment of Cholera. The Journal of Infectious Diseases 219:786–794.

51. Barman RK, Chakrabarti AK, Dutta S. 2022. Screening of Potential Vibrio cholerae Bacteriophages for Cholera Therapy: A Comparative Genomic Approach. Front Microbiol 13:803933.

52. Yang Y, Dufault-Thompson K, Yan W, Cai T, Xie L, Jiang X. 2024. Large-scale genomic survey with deep learning-based method reveals strain-level phage specificity determinants. GigaScience 13:giae017.

53. Kitts PA, Church DM, Thibaud-Nissen F, Choi J, Hem V, Sapojnikov V, Smith RG, Tatusova T, Xiang C, Zherikov A, DiCuccio M, Murphy TD, Pruitt KD, Kimchi A. 2016. Assembly: a resource for assembled genomes at NCBI. Nucleic Acids Res 44:D73–D80.

54. Seemann T. 2014. Prokka: rapid prokaryotic genome annotation. Bioinformatics 30:2068–2069.

55. Aramaki T, Blanc-Mathieu R, Endo H, Ohkubo K, Kanehisa M, Goto S, Ogata H. 2020. KofamKOALA: KEGG Ortholog assignment based on profile HMM and adaptive score threshold. Bioinformatics 36:2251–2252.

56. Jones P, Binns D, Chang H-Y, Fraser M, Li W, McAnulla C, McWilliam H, Maslen J, Mitchell A, Nuka G, Pesseat S, Quinn AF, Sangrador-Vegas A, Scheremetjew M, Yong S-Y, Lopez R, Hunter S. 2014. InterProScan 5: genome-scale protein function classification. Bioinformatics 30:1236–1240.

57. Dufault-Thompson K, Jiang X. 2024. Annotating microbial functions with ProkFunFind. mSystems 9:e00036–24.

58. Kanehisa M, Furumichi M, Tanabe M, Sato Y, Morishima K. 2017. KEGG: new perspectives on genomes, pathways, diseases and drugs. Nucleic Acids Res 45:D353–D361.

59. Katoh K. 2002. MAFFT: a novel method for rapid multiple sequence alignment based on fast Fourier transform. Nucleic Acids Research 30:3059–3066.

60. Suyama M, Torrents D, Bork P. 2006. PAL2NAL: robust conversion of protein sequence alignments into the corresponding codon alignments. Nucleic Acids Research 34:W609–W612.

61. Stamatakis A. 2006. RAxML-VI-HPC: maximum likelihood-based phylogenetic analyses with thousands of taxa and mixed models. Bioinformatics 22:2688–2690.

62. Lin Z, Akin H, Rao R, Hie B, Zhu Z, Lu W, Smetanin N, Verkuil R, Kabeli O, Shmueli Y, Dos Santos Costa A, Fazel-Zarandi M, Sercu T, Candido S, Rives A. 2023. Evolutionary-scale prediction of atomic-level protein structure with a language model. Science 379:1123–1130.

63. Gilchrist CLM, Chooi Y-H. 2021. clinker & clustermap.js: automatic generation of gene cluster comparison figures. Bioinformatics 37:2473–2475.

64. Schwechheimer C, Hebert K, Tripathi S, Singh PK, Floyd KA, Brown ER, Porcella ME, Osorio J, Kiblen JTM, Pagliai FA, Drescher K, Rubin SM, Yildiz FH. 2020. A tyrosine phosphoregulatory system controls exopolysaccharide biosynthesis and biofilm formation in Vibrio cholerae. PLoS Pathog 16:e1008745.

65. Price MN, Dehal PS, Arkin AP. 2010. FastTree 2 – Approximately Maximum-Likelihood Trees for Large Alignments. PLoS ONE 5:e9490.

66. Bayliss SC, Thorpe HA, Coyle NM, Sheppard SK, Feil EJ. 2019. PIRATE: A fast and scalable pangenomics toolbox for clustering diverged orthologues in bacteria. GigaScience 8:giz119.

67. Lees JA, Galardini M, Bentley SD, Weiser JN, Corander J. 2018. pyseer: a comprehensive tool for microbial pangenome-wide association studies. Bioinformatics 34:4310–4312.

68. Teufel F, Almagro Armenteros JJ, Johansen AR, Gíslason MH, Pihl SI, Tsirigos KD, Winther O, Brunak S, Von Heijne G, Nielsen H. 2022. SignalP 6.0 predicts all five types of signal peptides using protein language models. Nat Biotechnol 40:1023–1025.

69. Crooks GE, Hon G, Chandonia J-M, Brenner SE. 2004. WebLogo: A Sequence Logo Generator. Genome Res 14:1188–1190.

70. Rozewicki J, Li S, Amada KM, Standley DM, Katoh K. 2019. MAFFT-DASH: integrated protein sequence and structural alignment. Nucleic Acids Research gkz342.

71. Capella-Gutiérrez S, Silla-Martínez JM, Gabaldón T. 2009. trimAl: a tool for automated alignment trimming in large-scale phylogenetic analyses. Bioinformatics 25:1972–1973.

72. Letunic I, Bork P. 2024. Interactive Tree of Life (iTOL) v6: recent updates to the phylogenetic tree display and annotation tool. Nucleic Acids Research gkae268.

73. Guo J, Bolduc B, Zayed AA, Varsani A, Dominguez-Huerta G, Delmont TO, Pratama AA, Gazitúa MC, Vik D, Sullivan MB, Roux S. 2021. VirSorter2: a multi-classifier, expert-guided approach to detect diverse DNA and RNA viruses. Microbiome 9:37.

74. Bouras G, Nepal R, Houtak G, Psaltis AJ, Wormald P-J, Vreugde S. 2023. Pharokka: a fast scalable bacteriophage annotation tool. Bioinformatics 39:btac776.

