## Supplementary material for "Comprehensive Genomic and Evolutionary Analysis of Biofilm Matrix Clusters and Proteins in the *Vibrio* Genus": Supplementary_Figures.docx

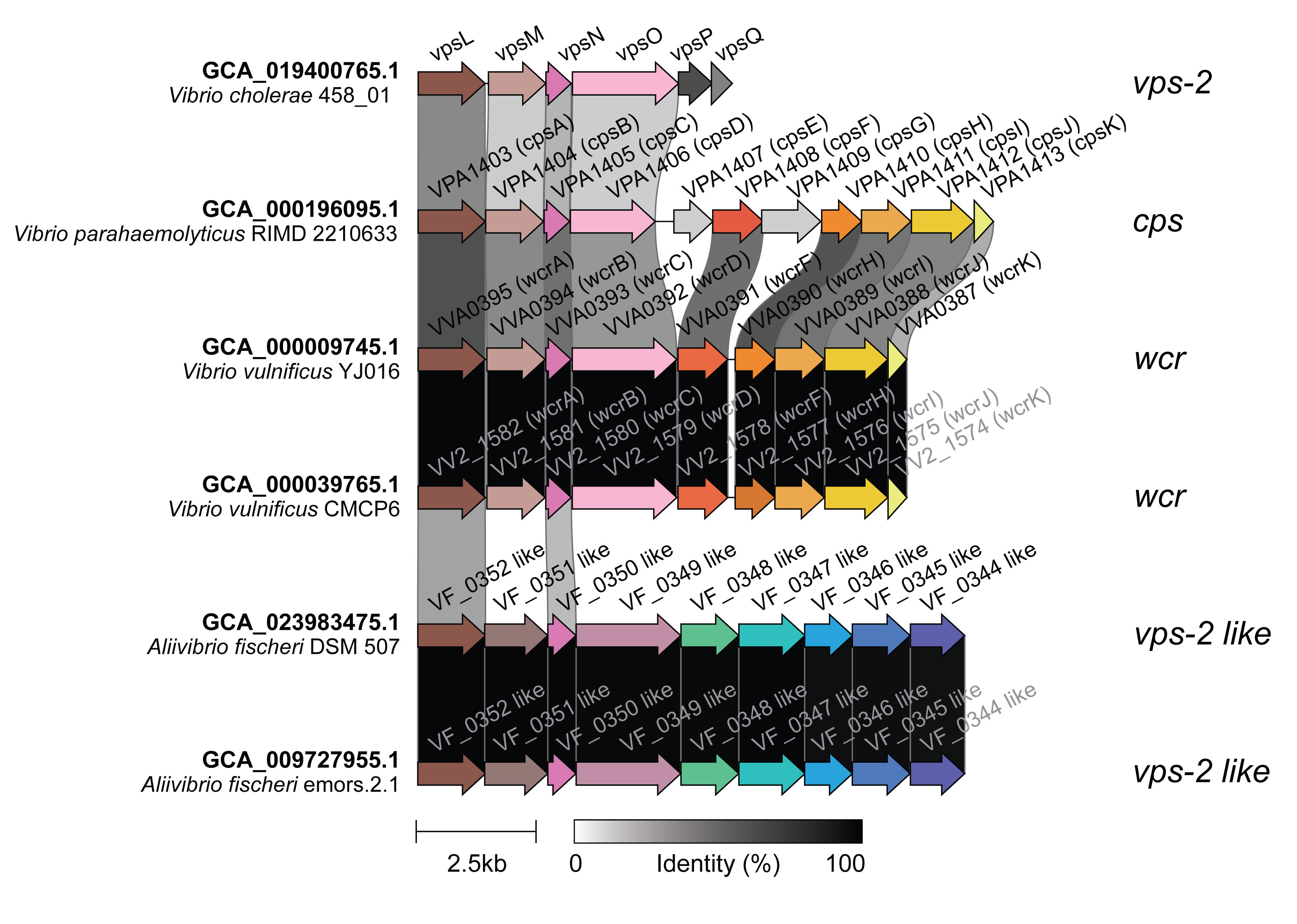

**Supplementary Figure 1.** Gene syntenies for *vps*-2 locus in *Vibrio cholerae,* *cps* locus in *Vibrio parahaemolyticus*, *wcr* loci in *Vibrio vulnificus*, and *vps*-2-like loci in *Alliivibrio fisheri* are depicted. Genes with more than 30% sequence similarity are color-coded. Link colors indicate sequence identities. The *cps*, *wcr*, and *vps*-2 loci all contain genes within their clusters that are similar to those found in the *vps*-2 cluster, particularly genes resembling *vpsLMNO*.

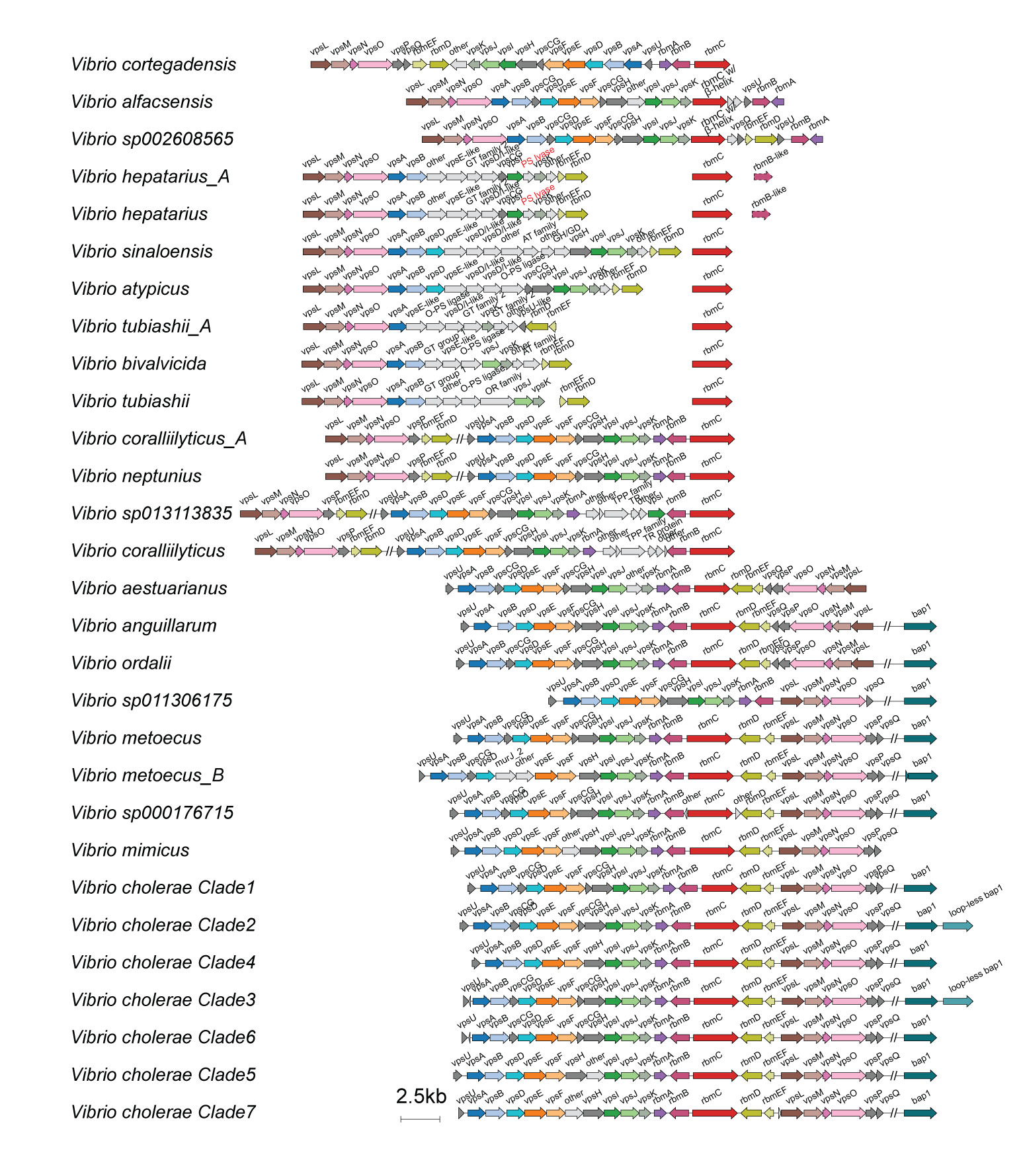

**Supplementary Figure 2.** Detailed gene syntenies for biofilm matrix clusters in 29 (sub)species with the same color palette as in Figure 1. GT: Glycosyltransferase; PS: Polysaccharide; O-PS: O-Antigen; AT: Acyltransferase; GH: Glycoside hydrolase; GD: Glycoside deacetylase; OR: Oxidoreductase; TPP: Thiamine pyrophosphate; TR protein: Transcriptional regulatory protein.

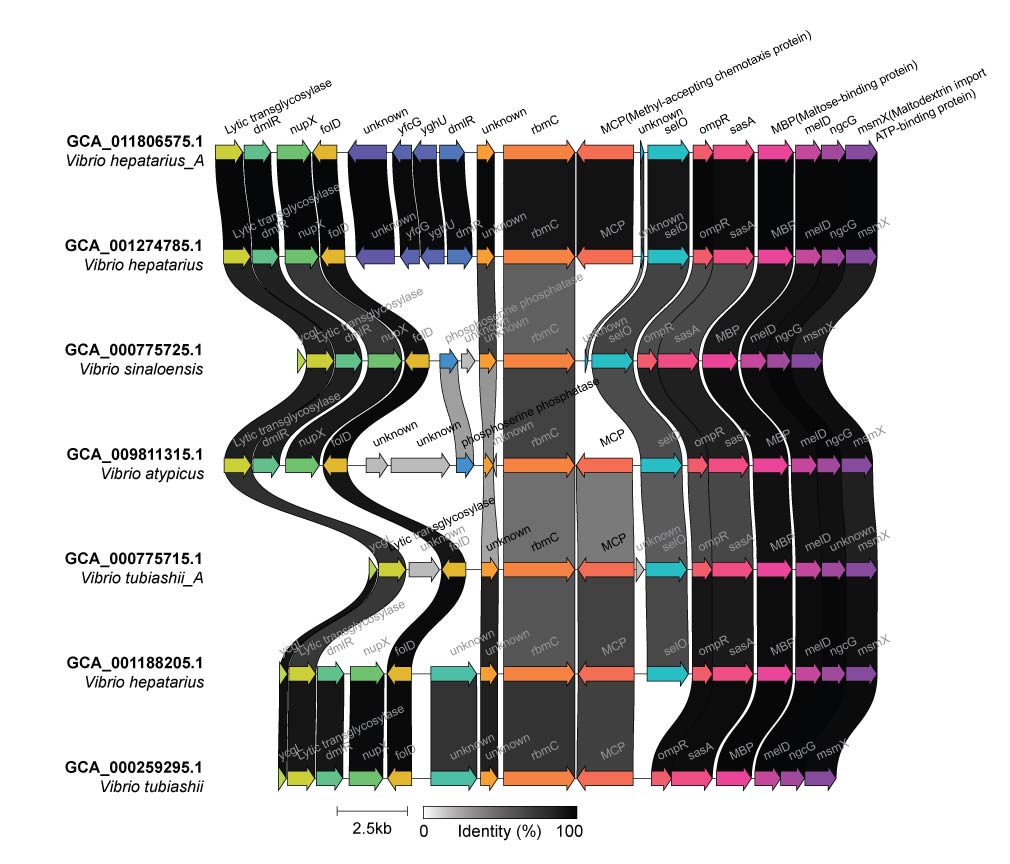

**Supplementary Figure 3.** Gene syntenies for *rbmC* genes and their neighboring genes in species with *rbmC* genes distant from the biofilm matrix cluster. Genes are color-coded by clusters sharing more than 80% sequence similarity, and link colors represent sequence identities. Detailed information is available in Supplementary Table 2.

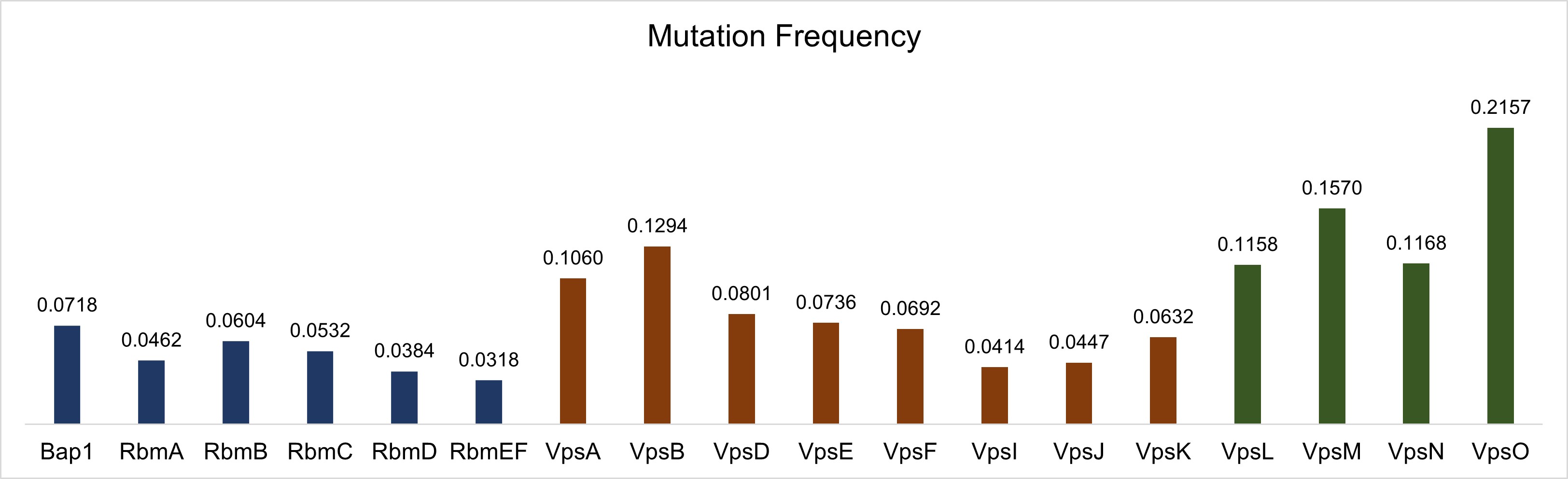

**Supplementary Figure 4.** Mutation frequencies calculated for 12 key *vps* genes and 6 *rbm* genes.

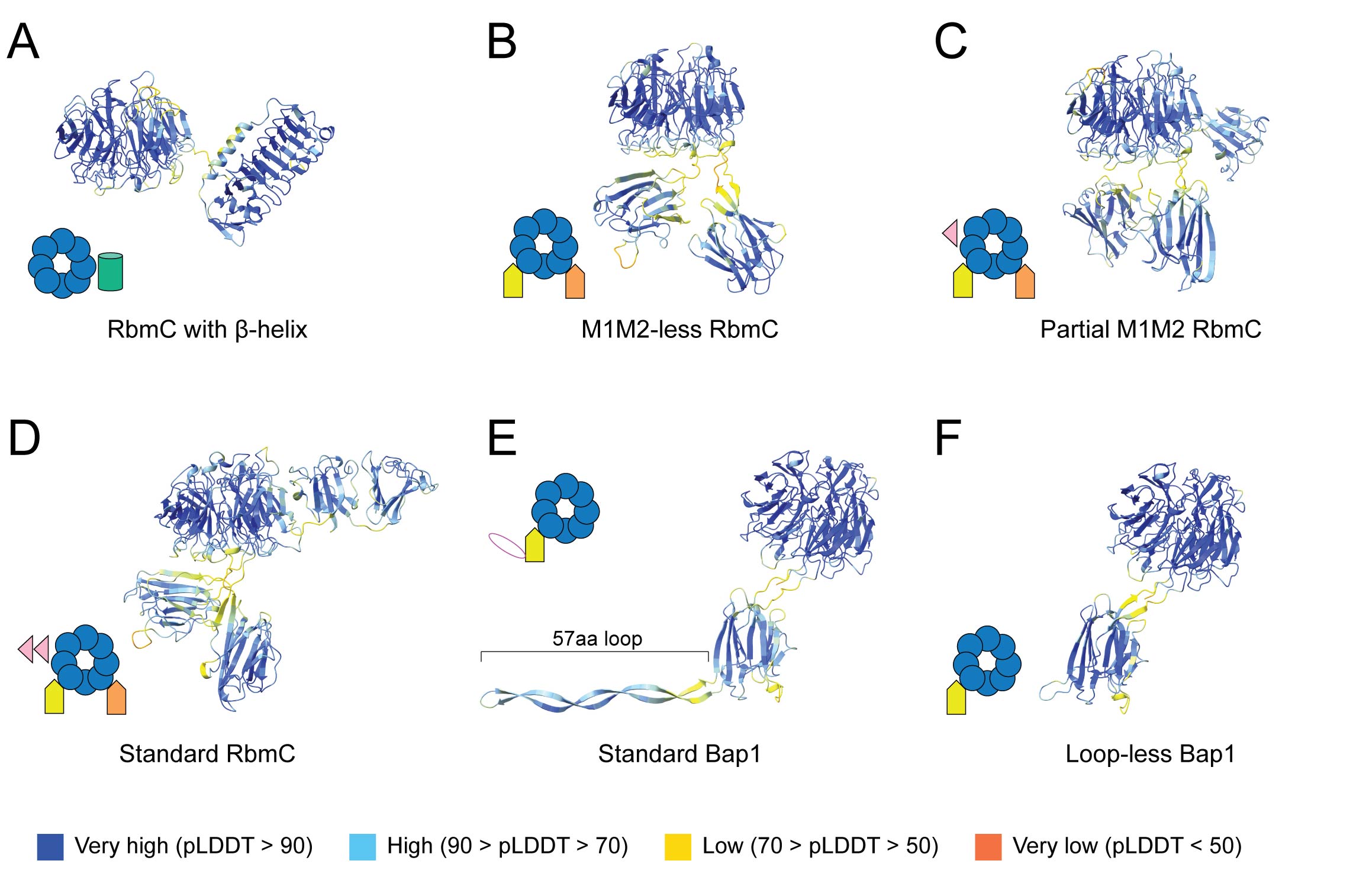

**Supplementary Figure 5.** Predicted structures by AlphaFold3 for proteins representing the six RbmC and Bap1 structural variants defined in this study. (A-F) Predicted structures of GCA_019670025.1_03371, GCA_003312035.1_01787, GCA_000259295.1_03774, GCA_019048845.1_03201, GCA_024746925.1_01708 and GCA_003716425.1_01353 representing proteins of RbmC with β-helix, M1M2-less RbmC, partial M1M2 RbmC, standard RbmC, standard Bap1 and loop-less Bap1, respectively.

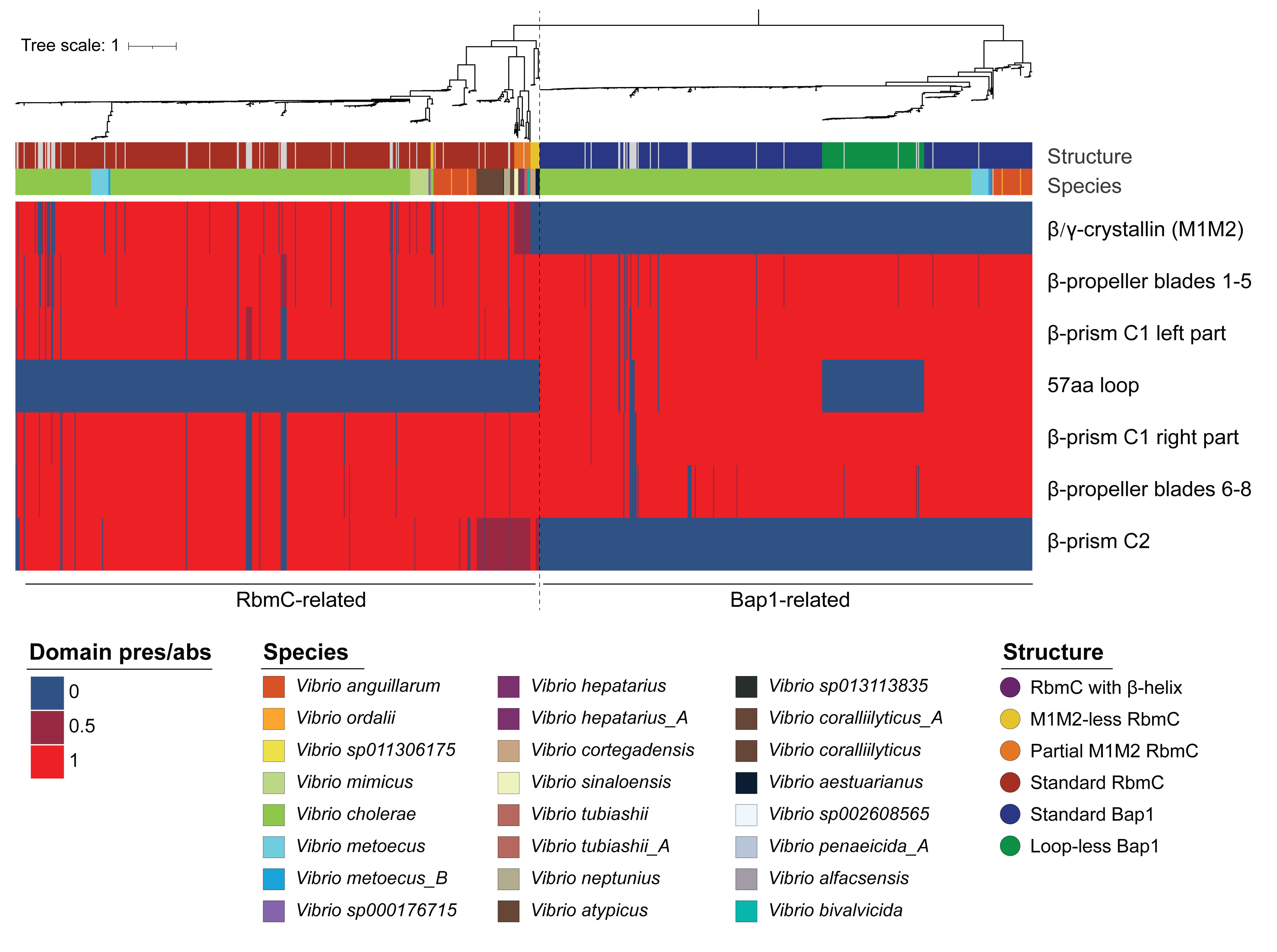

**Supplementary Figure 6.** Heatmap indicating domain presence and absence in 997 RbmC and Bap1 encoded genes. Rows represent domains. Columns represent genes and are mapped to the gene tree. The tips are annotated with the species of origin and structural types. Grey strips represent truncated proteins.

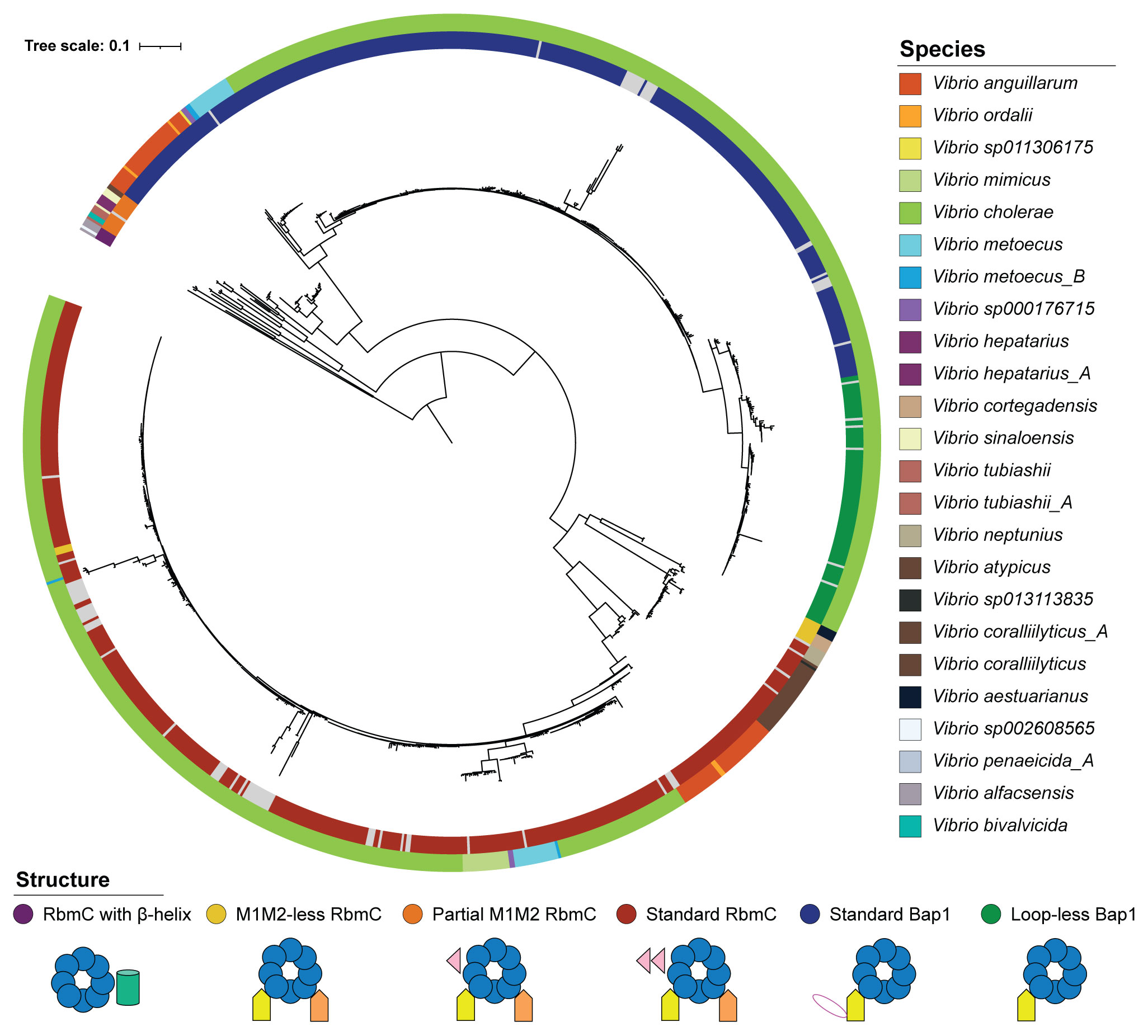

**Supplementary Figure 7.** The domain tree for 1,001 β-propeller domains of RbmC and Bap1 encoded sequences, rooted with RbmC with β-helix encoded genes. The outer circle indicates the species of origin, while the inner circle indicates the protein structural features. Grey strips represent truncated proteins.

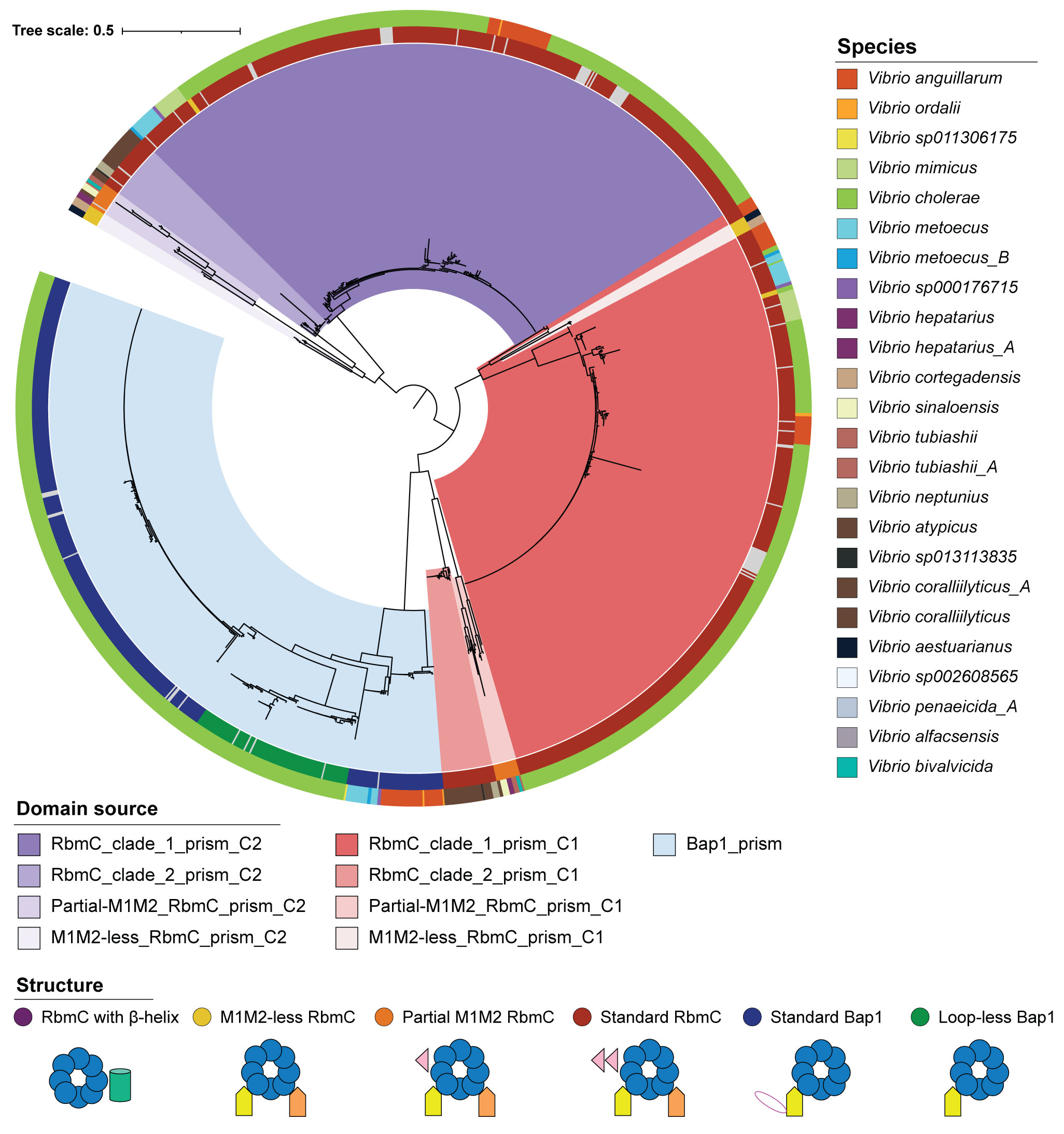

**Supplementary Figure 8.** The domain tree for 1,433 β-prism domains of RbmC and Bap1 encoded sequences, rooted at the midpoint. The outer circle indicates the species of origin, while the inner circle indicates the protein structural features. The color ranges indicate the domain source. Grey strips represent truncated proteins.

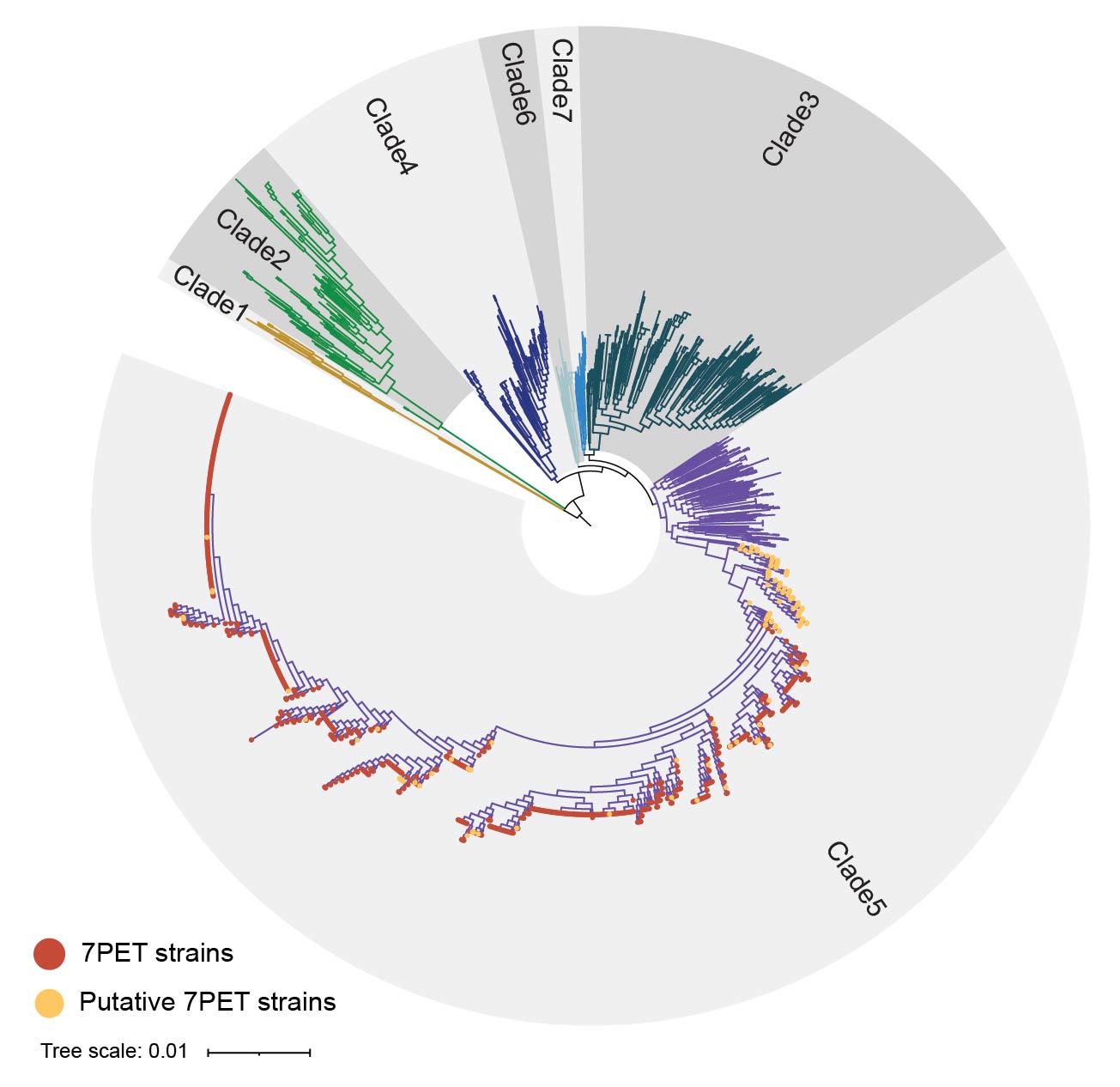

**Supplementary Figure 9.** The phylogenomic tree of *Vibrio cholerae* species (N=1,663), rooted at Clade 1. Strains identified as 7PET and putative 7PET lineages are marked with red and yellow dots at the tree tips, respectively.

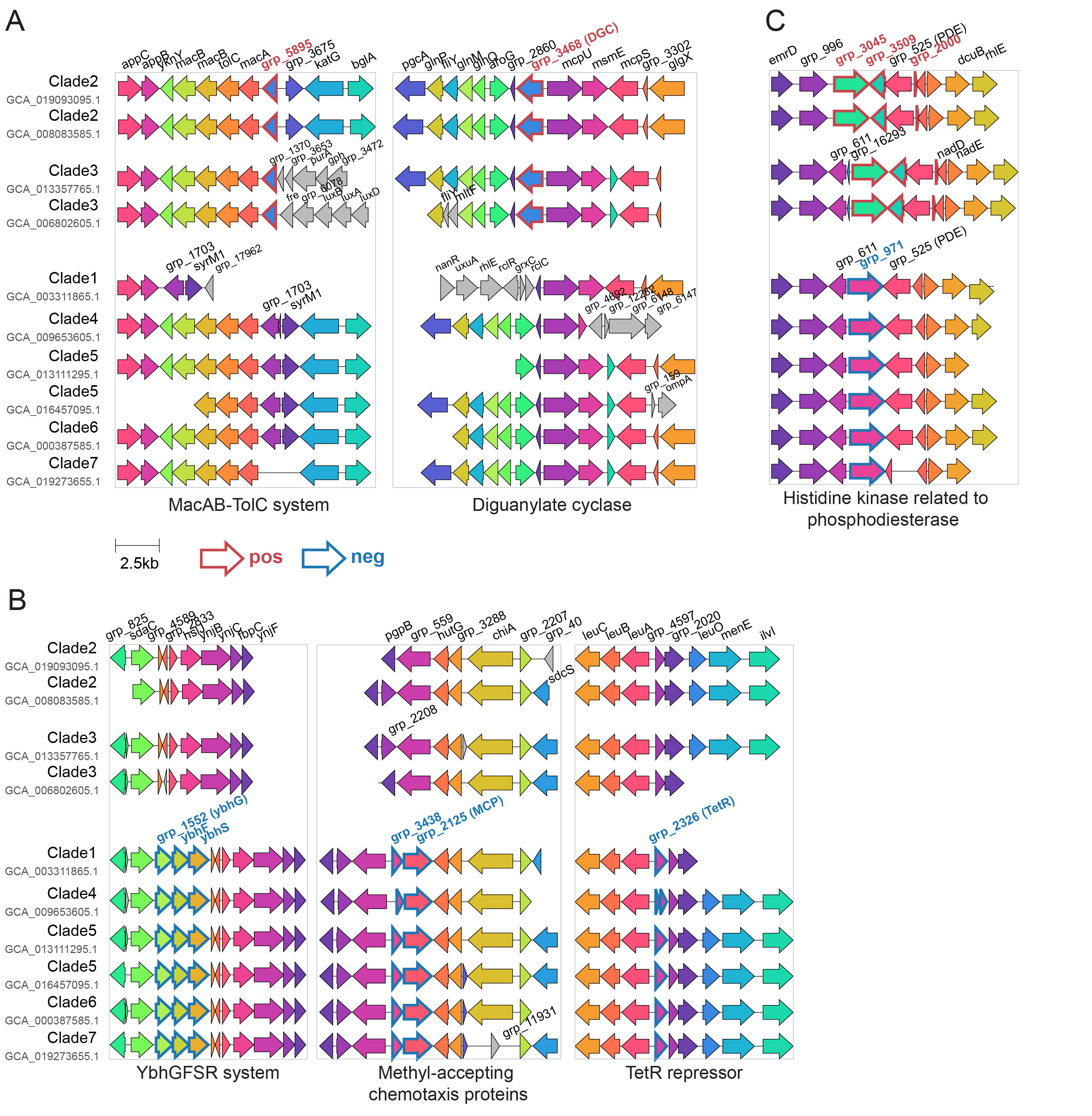

**Supplementary Figure 10.** Gene syntenies for associated gene groups in ten genomes selected from seven clades. They are highlighted by thicker red or blue borders to indicate their positive or negative associations, respectively. (A) Positively associated genes related to the MacAB-TolC system and diguanylate cyclase. (B) Negatively associated genes involved in YbhGFSR system, methyl-accepting chemotaxis proteins and TetR repressor. (C) Positively and negatively associated genes found in histidine kinases related to phosphodiesterase. Genes in the same boxes are colored by gene clusters sharing more than 80% sequence similarity. DGC: Diguanylate cyclase; PDE: Phosphodiesterase; MCP: Methyl-accepting chemotaxis protein; GTP: Guanosine-5’-triphosphate; GMP: Guanosine monophosphate.

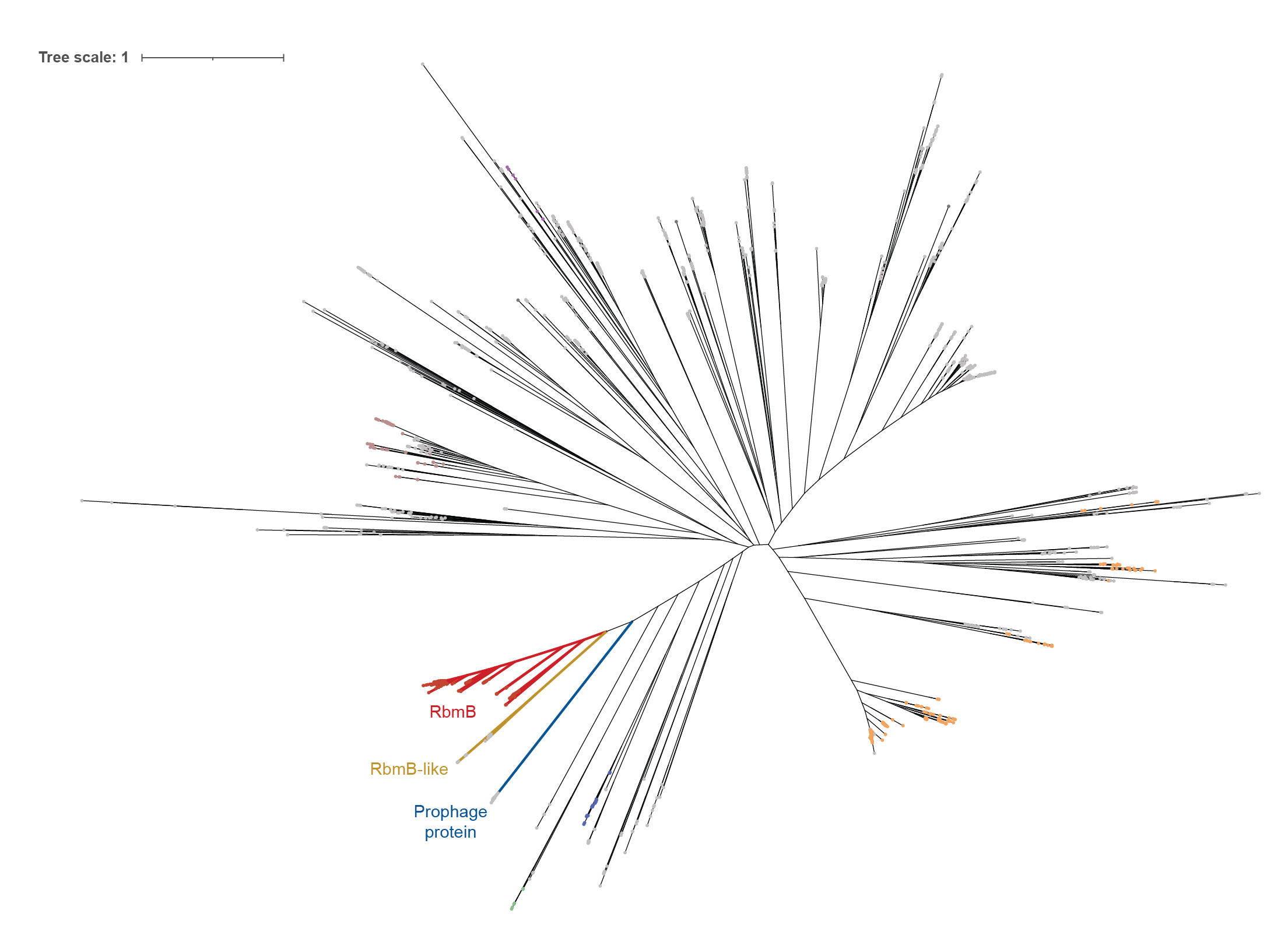

**Supplementary Figure 11.** Unrooted single-stranded right-handed β-helix domain containing gene tree. The branch colors highlight the clades for RbmB encoded genes (red), RbmB-like encoded genes (yellow) and prophage-related genes (blue).
