## Supplementary material for "Comprehensive Genomic and Evolutionary Analysis of Biofilm Matrix Clusters and Proteins in the *Vibrio* Genus": Supplementary_Information.docx

### **Supplementary Data**

**Data S1.** NCBI Assembly accessions and GTDB species for 6,121 Vibrio genomes.

**Data S2.** Biofilm matrix cluster and proteins annotation with ProkFunFind.

**Data S3.** RbmC and Bap1 protein classification table.

**Data S4.** RbmC and Bap1 protein sequences in FASTA format.

**Data S5.** 1,007 non-redundant predicted structures of RbmC and Bap1 proteins using ESMfold in PDB format.

**Data S6.** RbmB protein sequences in FASTA format.

**Data S7.** RbmA protein sequences in FASTA format.

**Data S8.** VpsE protein sequences in FASTA format.

**Data S9.** VpsF protein sequences in FASTA format.

**Data S10.** 216 representative genomes for *Vibrio* species.

**Data S11.** *V. cholerae* subspecies tree in NEWICK format.

**Data S12.** *Vibrio* species tree in NEWICK format.

**Data S13.** Gene groups detected in *V. cholerae* pangenome analysis using Roary.

**Data S14.** Signal peptide positions detected for RbmC and Bap1 using SignalP6.0.

**Data S15.** Single-stranded right-handed β-helix domain containing gene tree in NEWICK format.

**Data S16.** RbmC and Bap1 proteins’ β-propeller domain tree in NEWICK format.

**Data S17.** RbmC and Bap1 proteins’ β-prism domain tree in NEWICK format.

**Data S18.** Prophage regions detected in 1,803 genomes having single-stranded right-handed β-helix domain containing genes.
